# A bacterial effector directly targets Arabidopsis Argonaute 1 to suppress Pattern-triggered immunity and cause disease

**DOI:** 10.1101/215590

**Authors:** Odon Thiébeauld, Magali Charvin, Meenu Singla-Rastogi, Alvaro L Perez-Quintero, Fan Yang, Dominique Pontier, Pierre Barraud, Cécile Pouzet, Laure Bapaume, Delase Amesefe, Guangyong Li, Laurent Deslandes, Thierry Lagrange, James R. Alfano, Lionel Navarro

## Abstract

*Pseudomonas syringae* type III effectors were previously shown to suppress the Arabidopsis microRNA (miRNA) pathway through unknown mechanisms. Here, we first show that the HopT1-1 effector promotes bacterial growth by suppressing the Arabidopsis Argonaute 1 (AGO1)-dependent miRNA pathway. We further demonstrate that HopT1-1 interacts with Arabidopsis AGO1 through conserved glycine/tryptophan (GW) motifs, and in turn suppresses miRNA function. This process is not associated with a general decrease in miRNA accumulation. Instead, HopT1-1 reduces the level of AGO1-associated miRNAs in a GW-dependent manner. Therefore, HopT1-1 alters AGO1-miRISC activity, rather than miRNA biogenesis or stability. In addition, we show that the AGO1-binding platform of HopT1-1 is essential to suppress the production of reactive oxygen species (ROS) and of callose deposits during Pattern-triggered immunity (PTI). These data imply that the RNA silencing suppression activity of HopT1-1 is intimately coupled with its virulence function. Overall, these findings provide sound evidence that a bacterial effector has evolved to directly target a plant AGO protein to suppress PTI and cause disease.

## INTRODUCTION

Plants have evolved active immune responses to defend themselves against a wide range of pathogens. The plant immune system relies on the detection of pathogens by both surface-localized receptors and intracellular immune receptors. The recognition of Pathogen- or Microbe-Associated Molecular Patterns (PAMPs or MAMPs) is achieved by surface-localized Pattern-Recognition Receptors (PRRs) (Albert *et al*., 2020). Classical plant PRRs are composed of receptor-like kinases (RLKs) and receptor-like proteins (RLPs) (Albert *et al*., 2020). The most characterized plant PRRs are the leucine-rich repeat RLKs Flagellin Sensing 2 (FLS2) and EF-Tu Receptor (EFR), which recognize conserved epitopes from bacterial flagellin or elongation factor Tu, respectively (Gómez-Gómez and Boller, 2000; Zipfel *et al*., 2006). Upon ligand binding, these receptors initiate a complex phosphorylation cascade at the PRR complex that leads to early signaling events, which include the production of reactive oxygen species (ROS), the activation of mitogen-activated-protein-kinases (MAPKs), calcium signaling, and the differential expression of thousands of genes (Gómez-Gómez *et al*. 1999; Felix *et al*., 1999; Zipfel *et al*., 2004; Navarro *et al*., 2004; Zipfel *et al*., 2006; Couto and Zipfel, 2016; Köster *et al*., 2022). Later responses involve the biosynthesis of the phytohormone salicylic acid (SA) and cell wall modifications such as callose deposition, which ultimately culminate in Pattern-triggered immunity (PTI) (Hauck *et al*., 2003; Tsuda and Katagiri, 2010). Pathogens secrete virulence determinants, referred to as pathogen effectors, which can suppress PTI to cause disease (Jones and Dangl, 2006). The recognition of such divergent effector molecules is orchestrated by intracellular immune receptors, which belong to the nucleotide-binding domain (NBD), leucine-rich repeat (NLR) superfamily (Jones *et al*., 2016). This results in the formation of large protein complexes, referred to as resistosomes, which are critical to trigger NLR immune signaling (Huang *et al*., 2023). The transcriptome changes induced during such Effector-triggered immunity (ETI) immune responses significantly overlap with those activated during PTI (Tsuda and Katagiri, 2010; Navarro *et al*., 2004). It has also been reported that non-coding short interfering RNAs (siRNAs) and microRNAs (miRNAs) are differentially expressed during PTI and ETI, and a subset of these small RNAs can control gene expression as well as antimicrobial defense (Staiger *et al*., 2013; Jiang *et al*., 2023). This implies a key role of Post-Transcriptional Gene Silencing (PTGS) in the regulation of the plant immune system.

PTGS is an ancestral post-transcriptional gene regulatory process. The core mechanism of PTGS involves the production of double-stranded RNA (dsRNA) precursors, which are processed by DICER-LIKE (DCL) enzymes into 20-24 bp small RNA duplexes (Bologna and Voinnet, 2014). These small RNA duplexes associate with an Argonaute (AGO) protein, the central component of a multi-protein RNA-induced silencing complex (RISC) (Vaucheret, 2008). The guide small RNA further directs AGORISC to sequence complementary mRNA targets to trigger their post-transcriptional gene silencing. In plants, this phenomenon is manifested by endonucleolytic cleavage (so-called ‘slicing’) and/or translational inhibition of small RNA targets (Llave *et al*., 2002; Rhoades *et al*., 2002; Palatnik *et al*., 2003; Brodersen *et al*., 2008; Chen, 2004; Poulsen *et al*., 2013). *Arabidopsis thaliana* encodes 4 DCLs and 10 AGOs. DCL1 processes miRNA precursors with the help of other factors including the zinc-finger domain-containing protein SERRATE (SE) (Park *et al*., 2002; Finnegan *et al*., 2003; Kurihara and Watanabe, 2004; Lobbes *et al*., 2006). This reaction yields miRNA/miRNA* duplexes, where miRNA is the guide strand and miRNA* is the passenger strand. DCL2, DCL3 and DCL4 process endogenous and viral-derived dsRNAs into siRNA duplexes (Hajieghrari & Farrokhi, 2022). A significant proportion of dsRNAs is produced by RNA-dependent RNA polymerases (RDRs) that convert single-stranded RNAs (ssRNAs) into dsRNAs. RDR6, which is one of the six Arabidopsis RDRs, produces dsRNAs from viral transcripts, transposable elements, as well as from endogenous transcripts (Mourrain *et al*., 2000; Dalmay *et al*., 2000; Allen *et al*., 2005; Fei *et al*., 2013; Nuthikattu *et al*., 2013). In the latter pathway, the siRNAs are generated from the combined action of primary siRNA/miRNA-directed transcript targeting and of RDR6 activity, in conjunction with the RNA-binding protein SUPRESSOR OF GENE SILENCING 3 (SGS3), thereby resulting in the production of dsRNAs that are processed by DCL4 into 21nt phased secondary siRNAs, which are referred to as “phasiRNAs” (Allen *et al*., 2005; Xie *et al*., 2005; Fei *et al*., 2013, Arribas-Hernández *et al*., 2016; Mourrain *et al*., 2000; Béclin *et al*., 2002; Peragine *et al*., 2004). AGO1 is a major PTGS factor, which is loaded with miRNAs, (pha)siRNAs or viral-derived siRNAs and plays an important role in plant development (Bohmert *et al*., 1998; Fagard *et al*., 2000), antiviral defense (Morel *et al*., 2002) as well as bacterial PAMP-induced gene induction and callose deposition (Li *et al*., 2010; Wilkinson *et al*., 2023). Arabidopsis AGO1 can also positively or negatively regulate disease resistance against various fungal pathogens (Weiberg *et al*., 2013; Wang *et al*., 2016; Dunker *et al*., 2020; Cao *et al*., 2016; Zhao *et al*., 2023). Besides its role in PTGS, AGO1 also binds to the chromatin of active genes to promote their transcription by recruiting sRNAs and SWI/SNF chromatin remodeling complexes (Liu *et al*., 2018). AGO2 not only plays a critical role in antiviral silencing but also in antibacterial resistance mediated by the NLR RPS2 (Carbonell *et al*., 2012; Harvey *et al*., 2011; Jaubert *et al*., 2011; Zhang *et al*., 2011). AGO4 has also been reported to modulate antiviral silencing as well as antibacterial defense (Agorio and Vera, 2007; Brosseau *et al*., 2016).

Small non-coding RNAs have been implicated in various biological processes and play a key role in controlling plant-pathogen interactions. In the context of plant-viral interactions, viral-derived siRNAs repress translation, replication or accumulation of viral RNAs, thereby inhibiting viral replication (Hamilton and Baulcombe, 1999; Lopez-Gomollon & Baulcombe, 2022). In addition, plant miRNAs and siRNAs can modulate resistance against bacterial, fungal and oomycete phytopathogens by targeting either positive or negative regulators of PTI and/or ETI (Jiang *et al*., 2023). This phenomenon has been well characterized in the context of plant-bacterial interactions. As examples, miR393, miR160 and miR167 are PAMP-induced miRNAs that are loaded into Arabidopsis AGO1 to negatively regulate auxin signaling during PTI (Navarro *et al*., 2006; Fahlgren *et al*., 2007; Li *et al*., 2010), while miR393b* is enriched in Arabidopsis AGO2 during ETI, and targets a negative regulator of defense that acts downstream of RPS2 (Zhang *et al*., 2011). Furthermore, several endogenous siRNAs were found to be induced in response to a *P. syringae* strain carrying *AvrRpt2* and specifically required for RPS2-mediated resistance (Katiyar-Agarwal *et al*., 2006; 2007). It has also been reported that filamentous phytopathogens differentially regulate functionally relevant small RNAs during infection. For example, soybean miR393 is induced in response to *Phytophthora sojae* and positively regulates resistance against this oomycete pathogen (Wong *et al*., 2014). Besides their role in fine-tuning PTI and ETI responses, there is also growing evidence showing that some plant siRNAs and miRNAs are transferred from plant cells towards filamentous pathogens to trigger PTGS of essential and/or virulence factors (Wang *et al*., 2016; Cai *et al*., 2018; Hou *et al*., 2019; He *et al*., 2021, Qin *et al*., 2023).

Given that small non-coding RNAs play a major role in regulating plant immune responses as well as in targeting and silencing viral, fungal and oomycetal transcripts, it is not surprising that many pathogens have evolved PTGS suppression mechanisms to cause disease. This phenomenon has been extensively characterized in plant-viral interactions and we know now that most plant RNA viruses encode Viral Suppressors of RNA silencing (VSRs) (Lopez-Gomollon & Baulcombe, 2022). These proteins suppress different steps of PTGS and AGO1 has emerged as a critical VSR target (Zhang *et al.,* 2006; Derrien *et al*., 2012; Azevedo *et al*., 2010; Giner *et al*., 2010). Interestingly, RNA silencing suppressors were also reported from bacterial, oomycetal and fungal phytopathogens (Navarro *et al*., 2008; Qiao *et al*., 2013, Qiao *et al*., 2015; Hou *et al*., 2019; Yin *et al*., 2019, Zhu *et al*., 2022, Gui *et al*., 2022). In particular, we found that growth of a type III secretion defective mutant of *Pto* DC3000 and of non-adapted bacteria was significantly enhanced in Arabidopsis mutants that are impaired in miRNA biogenesis (Navarro *et al*., 2008). These results provide genetic evidence that the Arabidopsis miRNA pathway is essential for PTI and non-host resistance, and suggest that *Pto* DC3000 effectors must have evolved to suppress this small RNA pathway to enable disease. Accordingly, we have identified a series of Bacterial Suppressors of RNA silencing (BSRs) from this bacterium, that inhibit all the steps of the Arabidopsis miRNA pathway (Navarro *et al*., 2008). However, it remains unknown whether such BSRs could directly interact with components of the RNA silencing machinery and alter their activity as part of their virulence functions.

Here, we first found that the type III secreted Hrp outer protein T1-1 (HopT1-1) is a critical virulence determinant of *Pto* DC3000 that promotes pathogenesis by suppressing the AGO1-dependent miRNA pathway. We further show that HopT1-1 can interact with Arabidopsis AGO1 through two conserved GW motifs, which represent AGO-binding platforms previously found in some metazoan and plant AGO cofactors (Till *et al*., 2007; El-Shami *et al*., 2007; Azevedo *et al*., 2011). Importantly, we found that HopT1-1 does not alter the biogenesis nor the stability of miRNAs. Instead, it reduces the level of AGO1-bound miRNAs in a GW-dependent manner. This implies that HopT1-1 must act at the level of AGO1-miRISC activity rather than on miRNA biogenesis or stability. Finally, we found that the AGO-binding platform of HopT1-1 is essential for its ability to suppress PTI. These data indicate that the silencing suppression activity of HopT1-1 is coupled to its virulence function and relies, at least in part, on the targeting of Arabidopsis AGO1. Overall, this study provides sound evidence that a bacterial effector has evolved to directly target an AGO protein to suppress PTI and cause disease.

## RESULTS

### HopT1-1 is a key pathogenicity determinant that promotes growth of *Pto* DC3000 by suppressing the Arabidopsis AGO1-dependent miRNA pathway

HopT1-1 is an experimentally validated type III secreted protein expressed from the pDC3000A plasmid of *Pto* DC3000 (Guo *et al*., 2005). Although HopT1-1 was previously shown to suppress the transcriptional activation of a PAMP-responsive gene (Li *et al*., 2005), there is so far no evidence indicating a role for this effector in promoting bacterial multiplication *in planta*. To test this possibility, we first generated a *Pto* DC3000 mutant strain deleted of *hopT1-1*, hereafter referred to as *Pto* Δ*hopT1-1*, and assessed the ability of this strain to multiply in leaves of the Arabidopsis Col-0 reference accession (wild type plants, WT). Upon dip-inoculation of WT plants, we found that the *Pto* Δ*hopT1-1* mutant strain exhibited ∼ 10 times lower bacterial titer at 3 days post-inoculation (dpi) compared to the WT *Pto* DC3000 strain (Fig. 1 and Supplemental Fig. 1). This result indicates that HopT1-1 is a functionally relevant effector of *Pto* DC3000 that contributes to bacterial growth *in planta*.

**Fig. 1.**
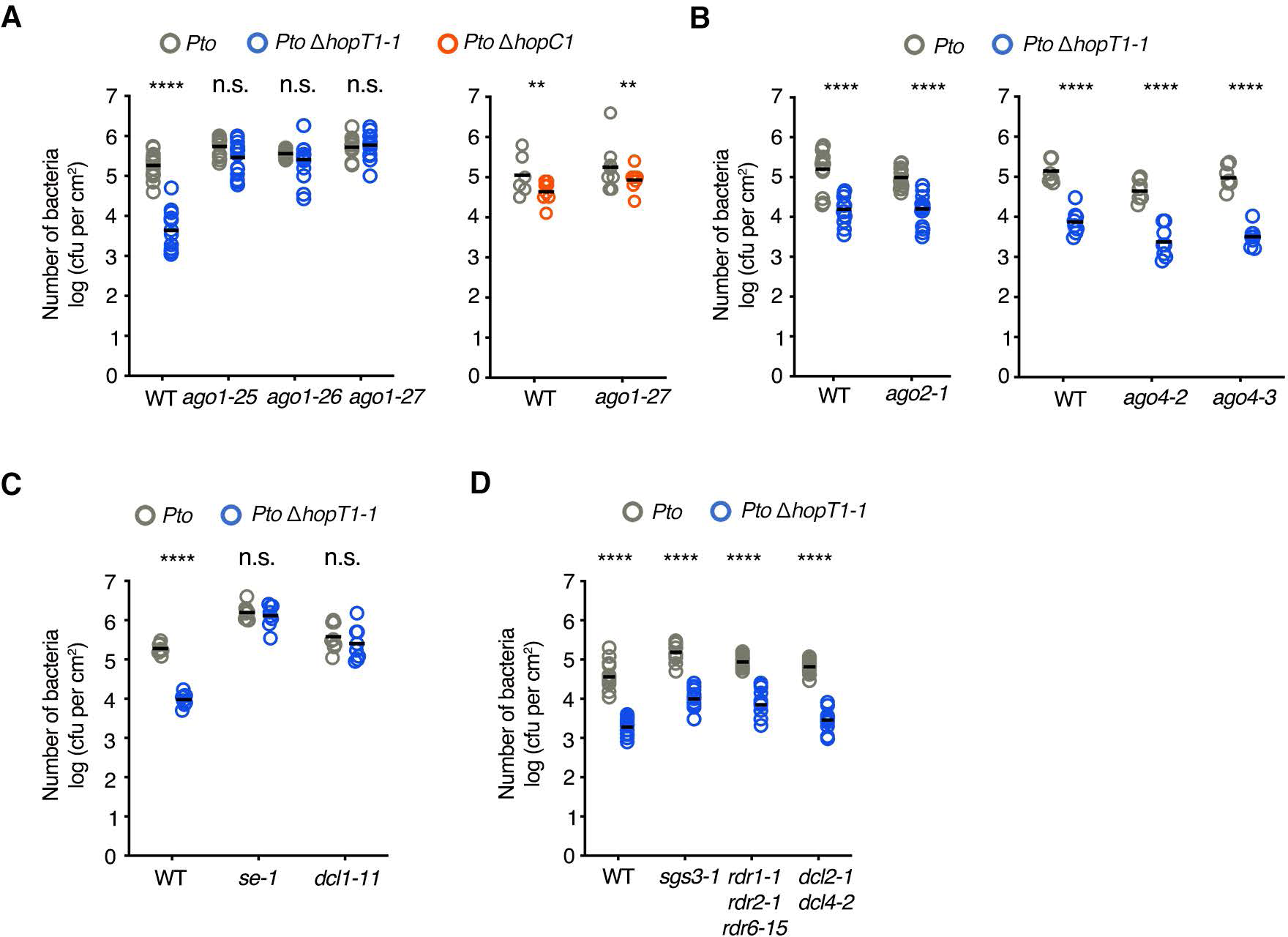
The growth defect of the *hopT1-1*-deleted strain of *Pto* DC3000 is specifically rescued in Arabidopsis miRNA-defective mutants. Five-week-old Col-0 Arabidopsis (WT) plants and indicated genotypes were dip-inoculated with bacterial strains *Pto* DC3000 (*Pto*) (grey dots) and *Pto ΔhopT1-1* (blue dots) or *Pto ΔhopC1* (orange dots) at a concentration of 10^8^ cfu/mL. At three days postinoculation (3 dpi), leaves from three plants were collected and bacterial titers were further monitored. Each dot represents a number of bacteria as log (cfu per cm^2^) and mean (n=8 or 16) is represented as horizontal line. Similar results were obtained in two to three independent experiments (results from biological replicates are provided in Fig. S1). **(A)** Left panel: growth of wild type *Pto* DC3000 strain (*Pto*) and of *hopT1-1* deleted bacterial strain (*Pto ΔhopT1-1)* in WT plants and in three different hypomorphic *ago1* mutants, namely *ago1-25*, *ago1-26* and *ago1-27*. Right panel: growth of wild type *Pto* DC3000 strain (*Pto*) and of *hopC1*-deleted bacterial strain (*Pto ΔhopC1)* in WT plants and in *ago1-27* mutant. **(B)** Growth of WT *Pto* and of *Pto ΔhopT1-1* in WT plants and in mutants defective in AGO2 (*ago2-1*) or in AGO4 (*ago4-2* and *ago4-3*). **(C)** Same as in (A-Left panel) but in miRNA biogenesis-defective mutants *se-1* and *dcl1-11*. **(D)** Same as in (B) but in siRNA biogenesis mutants: *sgs3-1, rdr1-1 rdr2-1 rdr6-15* and *dcl2-1 dcl4-2.* Statistical significance was assessed using the ANOVA test (n.s.: p-value > 0.05; *: p-value < 0.05; **: p-value < 0.01; ***: p-value < 0.001; ****: p-value < 0.0001).

HopT1-1 was previously shown to suppress AGO1-mediated miRNA- and siRNA-functions (Navarro *et al*., 2008), but the relevance of this interference in bacterial pathogenesis remains unknown. We took advantage of the *Pto* Δ*hopT1-1* mutant strain and examined whether its growth defect could be potentially rescued in *ago1* mutants. For this purpose, we dip-inoculated the *Pto* Δ*hopT1-1* strain on WT plants and on three hypomorphic *ago1* mutants, namely *ago1-25*, *ago1-26* and *ago1-27* (Morel *et al*., 2002), and subsequently monitored bacterial titers at 3 dpi. We also included in this assay the *ago2-1*, *ago4-2* and *ago4-3* mutants, as *AGO2* and *AGO4* were previously characterized in RPS2-mediated resistance and in antibacterial basal resistance, respectively (Zhang *et al*., 2011; Agorio and Vera, 2007). Importantly, the growth defect of *Pto* Δ*hopT1-1* was fully rescued in all *ago1* allelic mutants (Fig. 1A, Supplemental Fig. 1, A and B), while it remained unaltered in *ago2* and *ago4* mutants (Fig. 1B and Supplemental Fig. 1C). To further test whether the phenotype observed in *ago1* mutants was specific to the *Pto* Δ*hopT1-1* mutant strain, and not due to side effects caused by *ago1* developmental phenotypes, we repeated this assay with a *Pto* DC3000 mutant deleted of *hopC1* (*Pto* Δ*hopC1*). This effector partially contributes to *Pto* DC3000 multiplication in Arabidopsis WT leaves but does not interfere with miRNA action (Fig. 1A and Supplemental Fig. 1A; Navarro *et al*., 2008). Importantly, the partial *Pto* Δ*hopC1* growth defect observed in WT plants was not rescued in *ago1-27* plants (Fig. 1A and Supplemental Fig. 1A), indicating that the restoration of bacterial growth detected in *ago1* mutants was specific to the *Pto* Δ*hopT1-1* strain. Collectively, these results indicate that *AGO1* is a major genetic target of HopT1-1. They also suggest that AGO1 functions must be altered by HopT1-1 to promote growth of *Pto* DC3000 *in planta*.

Given that AGO1 is required for both miRNA- and siRNA-functions, we next assessed if each activity could be genetically targeted by HopT1-1 to promote growth of *Pto* DC3000 *in planta.* We first tested whether the *Pto* Δ*hopT1-1* growth defect could be rescued in *dcl1-11* and *se-1* mutants, which are both impaired in miRNA biogenesis (Lobbes *et al*., 2006; Zhang *et al*. 2008). Similar to the observation made in *ago1* mutant alleles, the *Pto* Δ*hopT1-1* growth defect was fully restored in these mutants (Fig. 1C, Supplemental Fig. 1, D and E). We then repeated the same assay in *sgs3-1*, *rdr1-1 rdr2-1 rdr6-15* triple and *dcl2-1 dcl4-2* double mutants, which are impaired in the biogenesis of endogenous siRNAs and viral-derived siRNAs (Mourrain *et al*., 2000; Dalmay *et al*., 2000; Xie *et al*., 2004; Deleris *et al*., 2006; Diaz-Pendon *et al*., 2007; Donaire *et al*., 2008), but did not find any growth restoration of the *Pto* Δ*hopT1-1* strain in these siRNA-defective mutants (Fig. 1D, Supplemental Fig. 1, F and G). Altogether, these results indicate that HopT1-1-triggered suppression of AGO1-mediated miRNA function is critical to promote growth of *Pto* DC3000 *in planta*.

### HopT1-1 interacts with Arabidopsis AGO1 through two conserved GW motifs

The above genetic data suggested that HopT1-1 could interact with Arabidopsis AGO1 to alter its miRNA-related function. The AGO-binding function of GW/WG platforms, present in some endogenous silencing factors as well as in some VSRs (El-Shami *et al*., 2007; Azevedo *et al*., 2011; Azevedo *et al*., 2010; Giner *et al*., 2010; Garcia *et al*., 2012; Aqil *et al*., 2013; Karran and Sansfaçon, 2014) prompted us to examine the protein sequence of HopT1-1 for the presence of such motifs. The *Pto* DC3000 HopT1-1 protein sequence contains three GW repeats at positions 79, 112 and 181 (Fig. 2A, Supplemental Fig. 2A). These motifs are referred to here as GW1, GW2 and GW3, respectively. It is noteworthy that the three motifs are present in the homologous sequence of *HopT1-2*, which is expressed from the *Pto* DC3000 chromosome, while only GW1 and GW2 are found in a truncated effector protein, a putative pseudogene that is closely related to HopT1-2 (Fig. 2, A and B; Wei *et al*., 2007). Importantly, the three GW motifs were found conserved across HopT1-1 homologous sequences derived from a subclade containing more distant bacteria, including the marine bacterium *Marinomonas mediterranea* MMB-1, but not in other subclades that are more phylogenetically distant (Fig. 2, A and B, Supplemental Fig. 2A). To test whether these GW motifs could be functional, we next generated tryptophan to phenylalanine substitutions (GW>GF) in each tryptophan residue of the GW motifs from the *Pto* DC3000 *HopT1-1* sequence. An *in*-*silico* analysis indicated that these point mutations have limited destabilization effects (Supplemental Fig. 2B). Furthermore, they do not alter the stability of HopT1-1 proteins when expressed *in planta* (Supplemental Fig. 3A). We further analyzed the ability of HopT1-1 WT and of the triple GW>GF mutant version, referred to as HopT1-1m3, to bind AGO1 *in planta*. To get a first insight into this possibility, we conducted bimolecular fluorescence complementation (BiFC) assays upon Agrobacterium-mediated transient transformation of *N. benthamiana* leaves, with constructs carrying the N-terminal fragment of the Yellow Fluorescent Protein (YFP) translationally fused to Arabidopsis AGO1, and the C-terminal fragment of the YFP fused with HopT1-1 or HopT1-1m3. In these experiments, we also used split-YFP fusions of HopC1 and of the silencing factor Silencing Defective 3 (SDE3) as negative controls. All these constructs were under the control of the moderately active *ubiquitin-10* promoter, which is suitable for transient expression of fluorescent-tagged proteins in *N. benthamiana* (Grefen *et al*., 2010). Confocal imaging revealed a clear fluorescence emission in epidermal cells co-expressing CYFP-HopT1-1/NYFP-AGO1 fusions, which was significantly different from the baseline fluorescent signal observed in cells co-expressing CYFP-HopC1/NYFP-AGO1 or CYFP-HopT1-1/NYFP-SDE3 fusions (Fig. 3A). These observations indicate that HopT1-1 is found in a close proximity to AGO1 *in planta*, which is not the case of SDE3 nor of HopC1, a *Pto* DC3000 effector that cannot suppress miRNA activity nor target AGO1 genetically (Fig. 1A and Supplemental Fig. 1A; Navarro *et al*., 2008). Importantly, we did not find any fluorescence emission upon coexpression of CYFP-HopT1-1m3/NYFP-AGO1 fusions (Fig. 3A), indicating that the GW motifs of HopT1-1 are essential to ensure its close proximity to AGO1 in plant cells. We next tested whether HopT1-1 could physically interact with AGO1 *in planta*. To this end, we used a non-invasive fluorescence imaging approach by conducting Föster resonance energy transfer-fluorescence lifetime imaging microscopy (FRET-FLIM) analyses after co-expressing a cyan fluorescent protein (CFP)-AGO1 (*35S_pro_:CFP-AGO1*) with either *35S_pro_:HopT1-1-YFP* or *35S_pro_:HopT1-1m3-YFP* constructs in *N. benthamiana* leaves. By doing so, we found a significant reduction in the average CFP lifetime of the donor CFP-AGO1 molecules in plant cells co-expressing CFP-AGO1/HopT1-1-YFP, compared with those co-expressing CFP-AGO1/HopT1-1-HA or expressing CFP-AGO1 alone (Fig. 3B, Supplemental Fig. 3, B and C), demonstrating a physical interaction between HopT1-1 and AGO1 *in planta*. Importantly, this proteinprotein interaction was fully dependent on the GW motifs of HopT1-1, because we did not detect any FRET in plant cells co-expressing CFP-AGO1 and HopT1-1m3-YFP (Fig. 3C, Supplemental Fig. 3B). Altogether, these data provide evidence that HopT1-1 interacts with AGO1 in plant cells, and that this process fully relies on its GW-dependent AGO-binding platform.

**Fig. 2.**
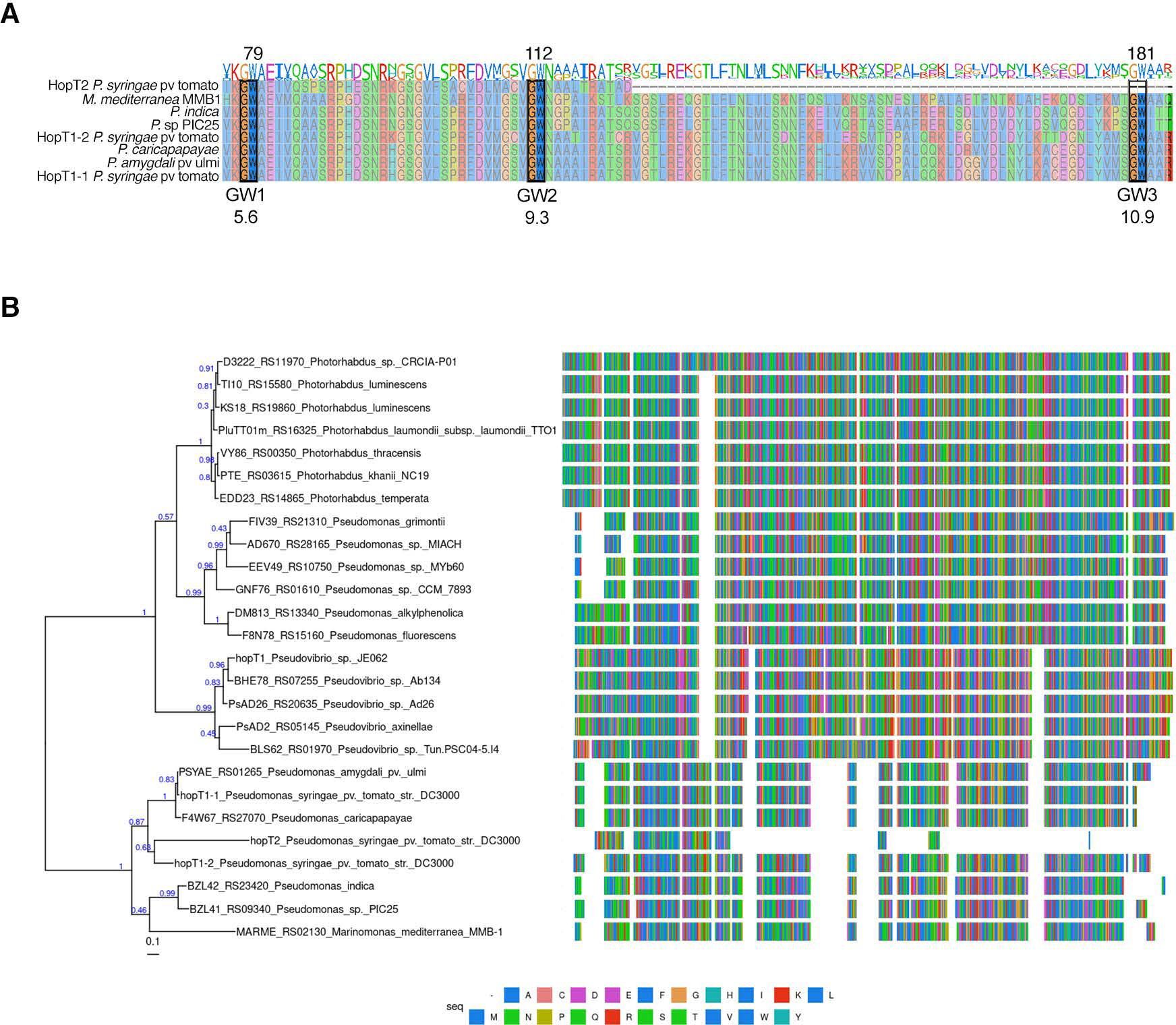
HopT1-1 possesses three GW motifs that are conserved among a subclade of bacteria. **(A)** Partial sequences (76-182 amino acids) of the *Pto* DC3000 HopT1 family were aligned using Muscle. These protein sequences possess three conserved GW motifs (named GW1, GW2 and GW3) highlighted in black boxes. The score of GW motifs prediction was retrieved by using the matrix AGO-planVir of the web portal http://150.254.123.165/whub/ (Zielezinski A. & Karlowski WM, 2015) and is indicated below each GW motif. **(B)** Maximum-likelihood phylogenetic tree (PhyML) based on a multiple alignment (right, obtained with Muscle) of amino-acid sequences of HopT1-1 orthologs. Sequences were obtained from defined orthology groups in OrthoDB v11 (Group 7800626at2). Numbers in blue indicate branch support as calculated by PhyML.

**Fig. 3.**
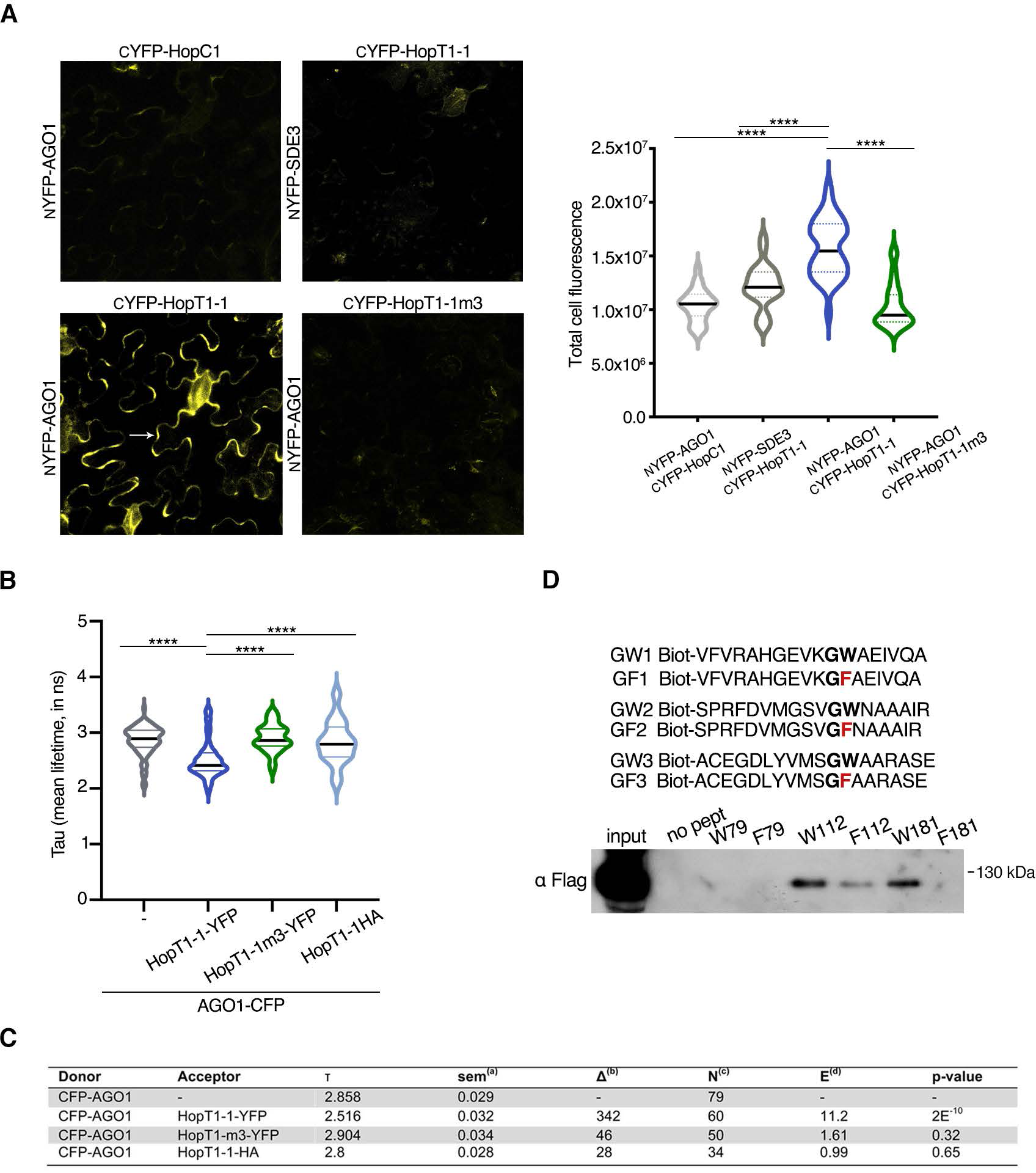
Two GW motifs of HopT1-1 are required for HopT1-1-AGO1 interaction. **(A)** BiFC assay in *N. benthamiana*. Four-week-old *N. benthamiana* were co-infiltrated with combinations of *Agrobacterium tumefaciens* strains carrying different constructs fused with N-terminal or C-terminal part of YFP. Interaction of NYFP-AGO1 with CYFP-HopT1-1 (left bottom) or with CYFP-HopT1-1m3 (right bottom) was tested. As negative controls, NYFP-AGO1 was co-infiltrated with CYFP-HopC1 (left top) and NYFP-SDE3 was co-infiltrated with CYFP-HopT1-1 (right top). YFP fluorescence was visualized by confocal microscopy three days post-infiltration (left panel) and quantification of the fluorescence signal for each picture taken (n=10) was performed using ImageJ software (right panel). Each dot represents the intensity of fluorescence signal in epidermal cells (white arrow) resulting from the interaction of each pair of combination as indicated in the dot plot. The signal in stomata cells is likely due to autofluorescence and should thus not be considered as positive BiFC signal. Two biological replicates are presented and the mean is represented as the horizontal line. Statistical significance was assessed using the ANOVA test (n.s.: p-value>0.05; ****: p-value<0.0001). **(B)** and **(C)** FRET-FLIM measurements showing that HopT1-1, but not HopT1-1m3, interacts with AGO1 in the cytoplasm of plant cells. Mean lifetime, τ, is in nanoseconds (ns). ^(a)^ For each cell analyzed, average fluorescence decay profiles measured in the cytoplasm were plotted and fitted with exponential function using a non-linear square estimation procedure, and the mean lifetime was calculated according to τ=∑ 〈_τ_τ_τ_^2^/∑ 〈_τ_τ_τ_ with I(t) = ∑ α_i_e^-t/τi^, standard error of the mean, ^(b)^ Δt=τ_D_-τ_DA_ (in ps), ^(c)^ total number of measured cell cytoplasm, and ^(d)^ % FRET efficiency: E = 1 – (τ_DA_/τ_D_). p-value of the difference between the donor lifetimes in the presence and in the absence of acceptor (Student’s *t* test) is indicated. **(D)** To assess which GW motif/s of HopT1-1 is/are required for the interaction with AGO1, synthetic biotinylated peptides containing the WT (GW) or the mutated version (GF) of each GW motif of HopT1-1 were mobilized on streptavidin magnetic beads. Specific peptide-loaded beads were further incubated with total protein extracted from Flag-AGO1 inflorescence. The presence of Flag-AGO1 in the total protein extract (input) and bound to the beads was assessed by immunoblotting. The peptides GW2 and GW3, but not GW1, can interact with Flag-AGO1. This interaction was partially or completely impaired in presence of GF2 and GF3, respectively.

To assess the contribution of each GW motif in HopT1-1-AGO1 interaction, we further chemically synthesized biotinylated peptides containing each GW motif surrounded by native amino acid residues. As negative controls, we synthesized mutated peptides with phenylalanine substitutions in the tryptophan of each GW motif (Fig. 3D). Equimolar amount of the peptides was bound to streptavidin columns (Supplemental Fig. 3D), and then incubated with inflorescence extracts from FLAG-AGO1 transgenic plants. After washing, eluted streptavidin-bound proteins were analyzed by Western blot using anti-FLAG antibody. Using this approach, we observed that the HopT1-1 GW2 and GW3 peptides, which exhibit the highest score for GW motif prediction using the Wsearch algorithm (Fig. 2A, Zielezinski and Karlowski, 2015), were both competent in binding FLAG-AGO1 proteins, while the HopT1-1 GW1 peptide was not (Fig. 3D). Furthermore, binding to FLAG-AGO1 was partially or completely lost upon incubation of inflorescence extracts from FLAG-AGO1 plants with the mutated HopT1-1 GF2 and GF3 peptides, respectively (Fig. 3D). Further Alphafold structural modeling of the HopT1-1 homologous sequences indicates that the GW1 residue is likely involved in HopT1-1 folding core formation and is buried in well-structured region, preventing its accessibility for interacting with AGO1. By contrast, this analysis shows that the functional GW2 and GW3 residues are partially exposed and adjacent to longer segments of potentially unstructured regions (pLDDT scores < 50) (Fig. 4A), supporting possible interactions with AGO1. These additional *in vitro* results therefore support our BiFC and FRET-FLIM data and indicate that HopT1-1 likely interacts with Arabidopsis AGO1 through two conserved motifs, namely GW2 and GW3.

**Fig. 4.**
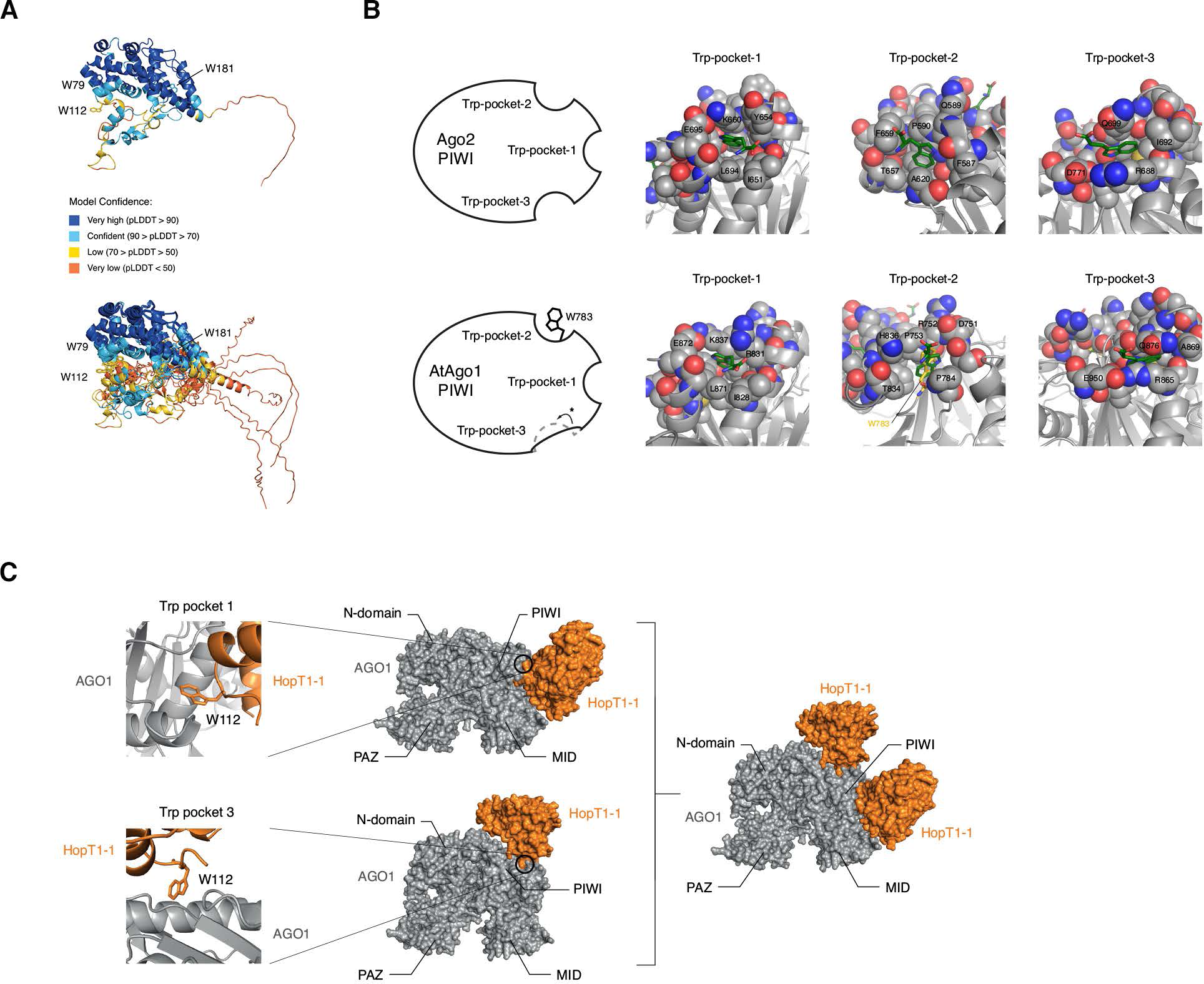
Structural modeling provides sensible assembly between the W112 residue from the GW2 motif of HopT1-1 and the two Trp-binding pockets of AGO1. **(A)** Upper panel: AlphaFold structural model of *Pto* DC3000 HopT1-1 (https://al-phafold.ebi.ac.uk/entry/Q88BP7). The structure is shown as a cartoon and the perresidue confidence score (pLDDT) is indicated: deep blue (very high, pLDDT > 90); light blue (confident, 90 > pLDDT > 70); yellow (low, 70 > pLDDT > 50); orange (very low, pLDDT < 50). Positions of W79, W112 and W181 are indicated. Bottom panel: Overlay of AlphaFold structural models of HopT1-1 protein family from the AlphaFold protein structure database (https://alphafold.ebi.ac.uk/ with the following accession numbers Q88BP7, F2JYU5, A0A0P9KI05, A0A2A2E7U8, A0A3M5KIL0, Q87WF7). The structures are shown as cartoons, and colored according to the per-residue confidence score (pLDDT). In HopT1-1 family, W79 is buried in a well-predicted and wellstructured region; and W112 and W181 are partially exposed/buried depending on the HopT1 models considered. **(B)** Upper panels: schematic (left panel) and close-up view (right panel) representations of the three Trp-pockets of human Ago2 PIWI domains according to the crystal structure of human Ago2. Bottom panels: schematic (left panel) and close-up view (right panel) representation of the putative three Trp-pockets of Arabidopsis AGO1 PIWI domain. The three Trp-pockets of Arabidopsis AGO1 from Alphafold model (https://alphafold.ebi.ac.uk/, accession number O04379) are pre-dicted based on human Ago2 Trp-pockets. Arabidopsis AGO1 Trp-pocket-1 is well conserved with human Ago2 Trp-pocket-1, while Arabidopsis AGO1 Trp-pocket-2 is already occupied *in cis* by a tryptophane residue (W783 in yellow), that is absent in human Ago2 Trp-pocket-2, making the interaction difficult with GW proteins. Arabidopsis AGO1 Trp-pocket-3 is only partially conserved with human Ago2 Trp-pocket-3, and appears as a partially closed pocket making AtAGO1 Trp-pocket-3 not available for interaction with GW proteins. Side-chain re-organization allows to re-open the pocket (* on the schematic representation of Arabidopsis AGO1 PIWI) to reach a human Ago2-like Trp-pocket-3, which could then interact with GW proteins. The bound tryptophanes are shown as green sticks and the residues forming the pockets are shown as spheres (grey for carbons, red for oxygen, blue for nitrogen and yellow for sulfur)**. (C)** HopT1-1 was manually docked through GW2 motif (W112) to Arabidopsis AGO1 Trp-pocket-1 (upper panel) and Trp-pocket-3 (lower panel). Left panel: close-up views of W112 overlayed with Arabidopsis AGO1 Trp-pockets. Right panel: overview of the HopT1-1 docking position on Arabidopsis AGO1 upon interaction of W112 HopT1-1 residue and Arabidopsis AGO1 Trp-pocket-1 and Trp-pocket-3. HopT1-1 and Arabidopsis AGO1 are shown in orange and grey, respectively. In this docking model, two HopT1-1 molecules have the possibility to be docked simultaneously to Arabidopsis AGO1 Trppocket-1 and Trp-pocket-3.

### Structural models constructed using the GW2 motif of HopT1-1, and the two potentially functional W-binding pockets of AGO1, unveil sensible assemblies

To get further insights into how HopT1-1 and AGO1 could interact with one another, we sought at modeling this interaction. For this end, we manually constructed models of HopT1-1 engaging interaction *via* either W112 from motif GW2 or W181 from motif GW3, with potentially functional tryptophan (W)-binding pockets of Arabidopsis AGO1. Because W-binding pockets have been extensively characterized in human AGO2 (Nakanishi, 2022), we first compared the Alphafold model of Arabidopsis AGO1 with the available structure of human AGO2 bound to tryptophan residues (pdb code 6cbd). Consistent with a previous report (Poulsen *et al*., 2013), this analysis suggests that the PIWI domain of Arabidopsis AGO1 retains at least one functional W-binding pocket (Trp-pocket-1), but probably also a second one (Trp-pocket-3) (Fig. 4B). Both W-binding pockets of the PIWI domain of Arabidopsis AGO1 are therefore likely available for interactions with the tryptophan residue of GW motifs. However, and as previously shown (Poulsen *et al*., 2013), the Arabidopsis AGO1 does not harbor the equivalent of the third W-binding pocket observed in the PIWI domain of human AGO2, because this pocket is filled *in cis* by a tryptophan of Arabidopsis AGO1, namely W783 (Fig. 4B).

We next conducted manual docking analyses between the W112 and W181 motifs of HopT1-1 and the two potentially functional W-binding pockets of the PIWI domain of Arabidopsis AGO1. Importantly, models constructed using W112 revealed sensible assemblies, in which only minor and limited structural conflicts occur between HopT1-1 and AGO1 (Fig. 4C). Models constructed using W181 generated less realistic assemblies, with extensive overlaps between AGO1 and HopT1-1 (Supplemental Fig. 3E). Interestingly, in the models generated using W112, the HopT1-1 molecules interacting with either AGO1 W-binding pocket-1 or W-binding pocket-3, occupy different regions of the three-dimensional space around AGO1, which allows for the possibility to accommodate two HopT1-1 molecules simultaneously, one in each W-binding pocket (Fig. 4C). These structural models illustrate how HopT1-1 could interact with the putative functional W-binding pockets of the PIWI domain of Arabidopsis AGO1, and put on display potential modes and stoichiometry of interaction. It is also noteworthy that it is not because we were unable to generate sensible assemblies *via* W181, when starting from the AlphaFold model of HopT1-1, that interaction of HopT1-1 with AGO1 is not occurring *via* this motif. Alteration of the W181 position and conformation compared to the AlphaFold model, could render the interaction of HopT1-1 with W-binding pockets of AGO1 structurally possible.

### HopT1-1 suppresses AGO1-mediated miRNA activity in a GW-dependent manner

To further test the requirement of the GW motifs of HopT1-1 in RNA silencing suppression, we have analyzed the ability of HopT1-1 and HopT1-1m3 to suppress miRNA activity *in vivo*. To this end, we transformed WT Arabidopsis with either a *35S*_pro_*::HopT1-1* or a *35S*_pro_*::HopT1-1m3* construct and further monitored through RNA sequencing (RNA-seq) the extent to which miRNA targets were differentially expressed in primary transformants. Principal component analysis (PCA) revealed that the biological replicates from the *35S*_pro_*::HopT1-1* plants were distant from those of WT and *35S*_pro_*::HopT1-1m3* plants, supporting an impact of the AGO-binding platform of HopT1-1 on the Arabidopsis transcriptome (Supplemental Fig. 4A). More specifically, when we analyzed the levels of experimentally validated miRNA targets in HopT1-1 transgenic plants compared to WT plants, we found that a large proportion of them were differentially expressed, with a majority being derepressed, as previously reported in *ago1-3* mutants (Arribas-Hernandez *et al*., 2016) (Fig. 5A). By contrast, none of these miRNA targets were differentially expressed in HopT1-1m3 transgenic plants compared to WT (Fig. 5A), supporting a key role of the AGO-binding platform of HopT1-1 in this process. To validate these findings, we further performed real-time quantitative PCR (RT-qPCR) analyses on a subset of miRNA targets, in individual primary transformants (n = 12 or 13 individuals) expressing comparable levels of *HopT1-1* and of *HopT1-1m3* (Supplemental Fig. 4, B and C). We found a significant enhanced mRNA accumulation of *SPL10* (miR156), *MYB33* (miR159), *ARF10* (miR160), *ARF17* (miR160), *PHB* (miR166) and *HAP2B* (miR169) in HopT1-1 lines compared to WT plants (Fig. 5B, Supplemental Fig. 4D). Derepression of *SPL10* (miR156), *ARF17* (miR160), *PHB* (miR166) and *HAP2B* (miR169) was also detected in hypomorphic *ago1-27* mutants (Fig. 5B), which served as positive controls in these RT-qPCR experiments. In addition, the levels of *AGO1*, *AGO2* and *DCL1* transcripts, which are controlled by miR168, miR403 and miR162/838, respectively (Xie *et al*., 2003; Vaucheret *et al*., 2004; Allen *et al*., 2005; Rajagopalan *et al*., 2006), were more elevated in HopT1-1 lines compared to WT plants (Fig. 5C, Supplemental Fig. 4E), and this effect was associated with an enhanced accumulation of cognate proteins in these transgenic lines as revealed by Western blot analyses (Fig. 5D, Supplemental Fig. 4F). An enhanced accumulation of *AGO1* and *AGO2* mRNAs, as well as of AGO2 and DCL1 proteins, was also observed in *ago1-27* compared to WT plants (Fig. 5C, 5D). However, we found that AGO1 protein levels remained low in *ago1-27* plants (Fig. 5D), which is probably due to the remaining negative regulation exerted by AGO10 over AGO1 protein accumulation in this *ago1* hypomorphic mutant (Mallory *et al*., 2009). Furthermore, mRNA and protein levels of AGO4, which is not targeted by miRNAs, were unchanged in HopT1-1 transgenic lines and in *ago1-27* plants compared to WT plants (Fig. 5, C and D), indicating a specific effect of HopT1-1 over miRNA targets. In addition, we found that all the above miRNA targets remained fully silenced in HopT1-1m3 plants (Fig. 5, B to D, Supplemental Fig. 4, D to F), implying a key role of the AGO-binding platform of HopT1-1 in their derepression. Collectively, these data provide evidence that HopT1-1 suppresses AGO1-dependent miRNA activity in a GW-dependent manner.

**Fig. 5.**
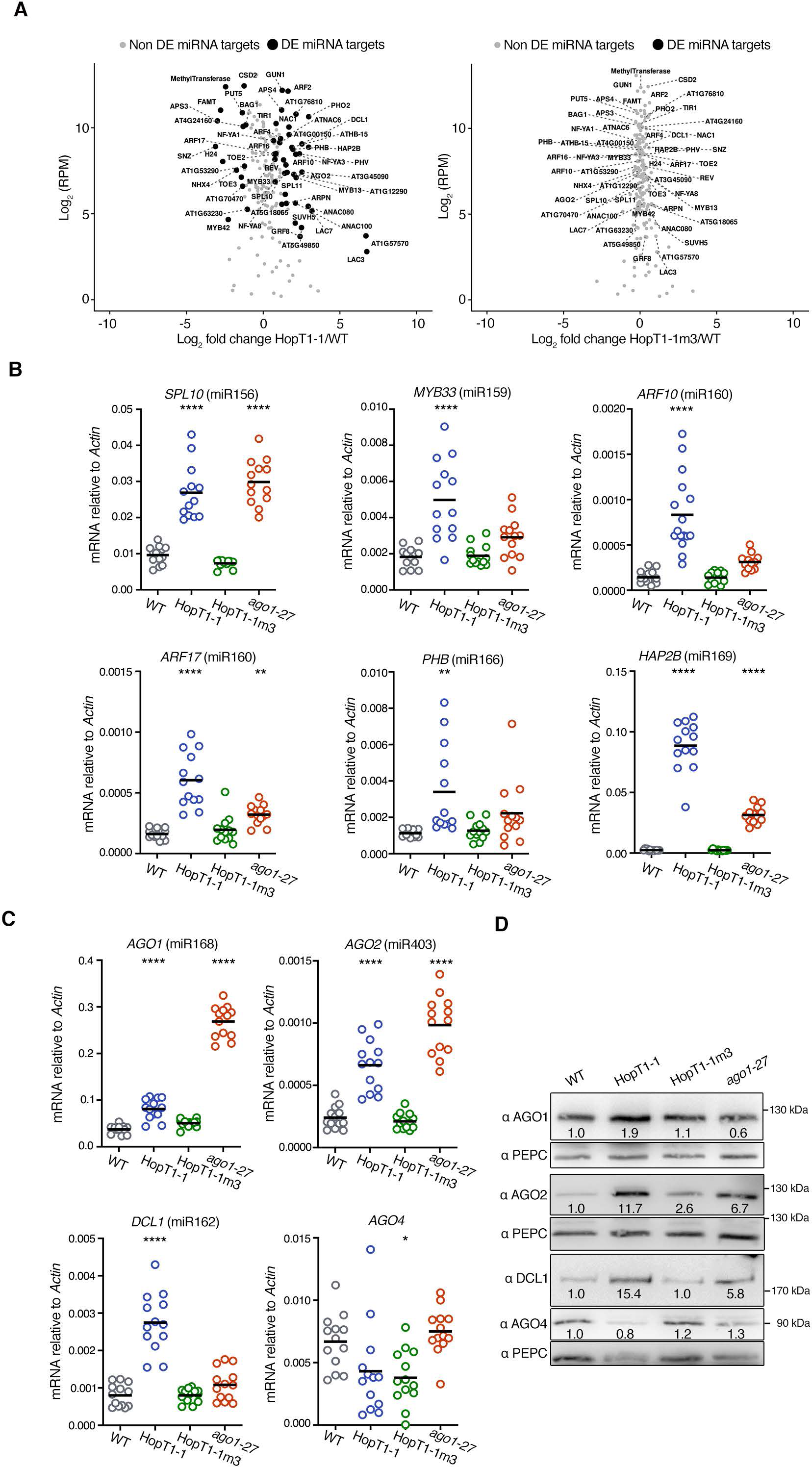
HopT1-1 suppresses AGO1-mediated miRNA function in a GW-dependent manner. **(A)** Volcano plots showing average reads per million (log_2_ RPM) for genes targeted by miRNAs *versus* log_2_ fold change when comparing HopT1-1 and WT RNA-seq libraries (left panel), and HopT1-1m3 vs WT RNA-seq libraries (right panel). DE genes (adjusted p-value <0.05) that are known microRNA targets are shown as black circles. Labeled points are DE microRNA targets in the HopT1-1/WT comparison. **(B)** Expression of endogenous miRNA targets was monitored by RT-qPCR analysis in the same HopT1-1 transgenic plants as well as in individual WT and *ago1-27* mutant plants. The miRNA that targets the analyzed transcript is indicated between brackets. *Actin* was used as a control. **(C)** Relative mRNA accumulation levels of RNA silencing factors targeted by miRNAs were monitored by RT-qPCR analysis in the same plants described in (B). AGO4 was used as an internal control, as this silencing factor is not targeted by any miRNA. *Actin* was used as a control. Statistical significance was assessed by comparing the mean (black bar) of each condition with the mean of WT condition, using one-way ANOVA analysis (*: p-value < 0.05; **: p-value < 0.01; ****: p-value < 0.0001). **(D)** Individuals of the plants depicted in (B) were pooled to monitor by immunoblotting the protein accumulation levels of the RNA silencing factors depicted in (C). PEPC protein accumulation level was used as loading control for each blot. Presence of DCL1 and AGO4 proteins was revealed using the same blot. Relative quantification of the protein accumulation using PEPC accumulation level was done using ImageJ software and is indicated below each condition. Another biological replicate is shown from other individual plants pooled in Fig. S4.

### HopT1-1 does not alter the biogenesis nor the stability of canonical miRNAs

We next assessed whether the ability of HopT1-1 to suppress miRNA function could be caused by an alteration in miRNA biogenesis and/or stability. For this purpose, we sequenced small RNAs from adult leaves of WT plants, *35S*_pro_::HopT1-1 and *35S*_pro_::HopT1-1m3 transgenic lines. We found that the overall size distribution and abundance of miRNAs were similar in *35S*_pro_::HopT1-1 compared to WT or *35S*_pro_::HopT1-1m3 plants (Fig. 6A). Accordingly, we did not detect statistically significant changes in the overall abundance of miRNAs in *35S*_pro_::HopT1-1 plants (Fig. 6B). These data indicate that HopT1-1 does not exhibit a general effect on miRNA biogenesis nor on miRNA stability. To verify these findings, we further monitored the accumulation of a subset of conserved miRNAs, namely miR156, miR159, miR160, miR166, miR167, miR168, miR393 and miR398, by low molecular weight northern blot analysis. Similarly, we did not detect consistent changes in the accumulation of these miRNAs across the different biological replicates in *35S*_pro_::HopT1-1 *versus* WT plants or *35S*_pro_::HopT1-1m3 plants (Fig. 6C, Supplemental Fig. 5A). Therefore, the *in planta* expression of HopT1-1 does not interfere with the accumulation of these mature miRNAs. Finally, we monitored the levels of primary miRNA transcripts in those plants by RT-qPCR analysis. We found that *pri-miR156a*, *pri-miR166a* and *pri-miR167a* were either unchanged or barely affected in *35S*_pro_::HopT1-1 plants compared to WT or *35S*_pro_::HopT1-1m3 plants (Supplemental Fig. 5, B and C). A mild increase in the accumulation of *pri-miR159b*and *pri-miR168a* was specifically detected in *35S*_pro_::HopT1-1 plants compared to WT or *35S*_pro_::HopT1-1m3 plants (Supplemental Fig. 5, B and C). However, these effects were much weaker than the ones typically detected in the *dcl1-11* and *se-1* mutants, which are impaired in miRNA biogenesis (Supplemental Fig. 5, B and C; Zhang *et al*., 2008; Yang *et al*., 2006; Lobbes *et al*., 2006). Altogether, these data indicate that the ability of HopT1-1 to suppress miRNA function is not due to a general inhibitory effect on miRNA biogenesis nor on miRNA stability.

**Fig. 6.**
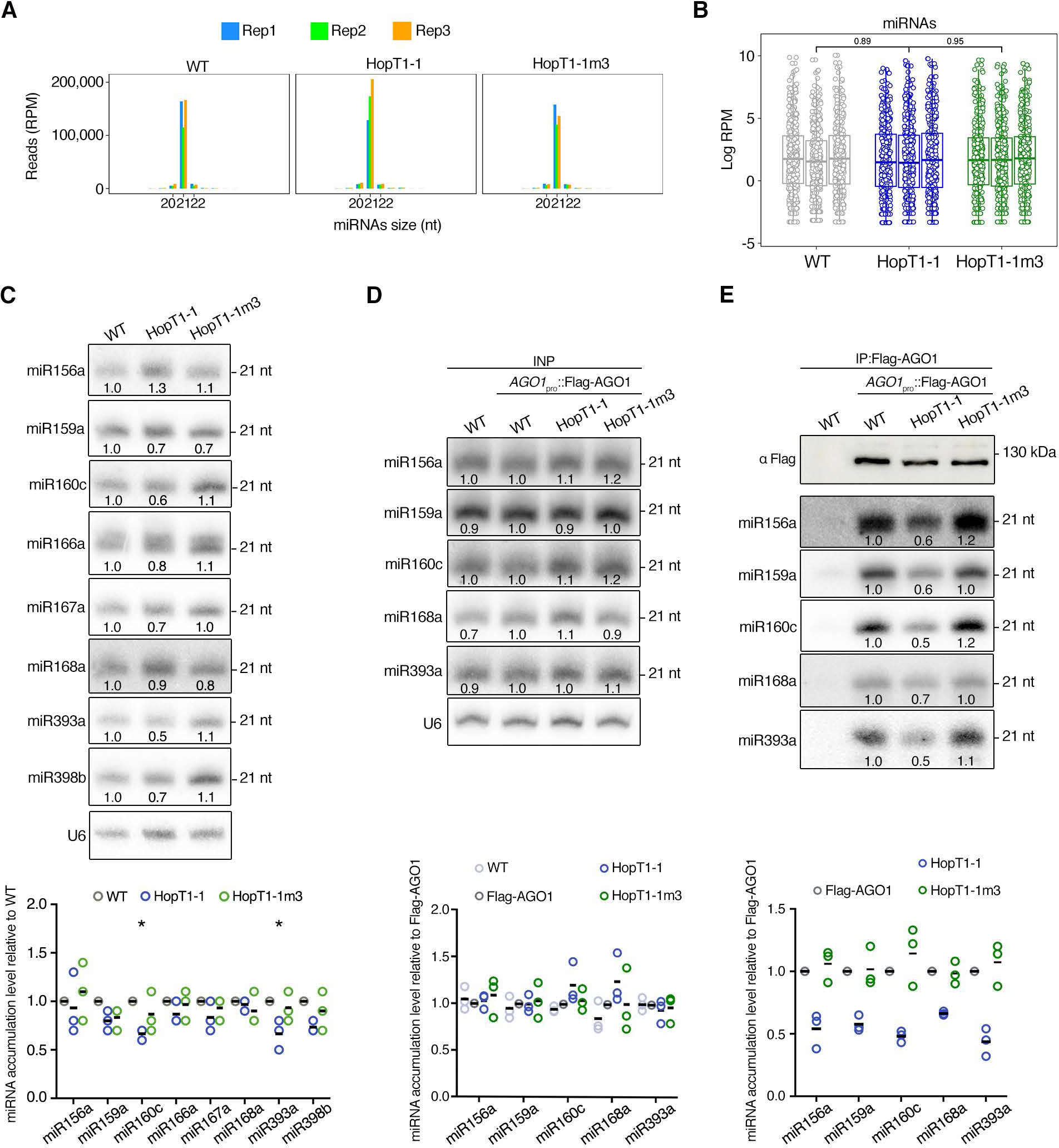
The AGO-binding platform of HopT1-1 does not alter the biogenesis/stability of miRNAs but interferes with the association of miRNAs with AGO1. **(A)** Total accumulation level (in reads per million, rpm) of all annotated (miRbase) Arabidopsis miRNAs according to their size (in nucleotides, nt) in WT plants or in the transgenic lines expressing HopT1-1 or HopT1-1m3. Three biological replicates are represented. **(B)** Accumulation level of each miRNA (log reads per million, rpm) annotated in miRbase in WT plants or in the transgenic lines expressing HopT1-1 or HopT1-1m3. Three biological replicates are represented. Numbers indicate p-value of a *t* test comparing treatments with the expression of each microRNA averaged across replicates. **(C)** Top panel: accumulation levels of endogenous miRNAs in pooled WT plants and in pooled primary transformants expressing HopT1-1 or HopT1-1m3 were assessed by northern blot. U6 was used as a loading control. Relative quantification of each miRNA accumulation using U6 accumulation level was done using ImageJ software and is indicated below each condition. Bottom panel: quantification of the miRNA accumulation level in HopT1-1 and in HopT1-1m3 plants relative to the miRNA accumulation in WT plants is shown for three biological replicates (Fig. S5A). **(D)** Inflorescences from WT plants or from *AGO1*_pro_::Flag-AGO1 transgenic line or from pooled primary transformants expressing HopT1-1 or HopT1-1m3 in *AGO1*_pro_::Flag-AGO1 background were collected, in order to analyze the accumulation level of canonical miRNAs by Northern blot (top panel). U6 was used as a loading control using U6 accumulation level was done using ImageJ software and is indicated below each condition. Bottom panel: quantification of the miRNA accumulation level in *AGO1*_pro_::Flag-AGO1 transgenic plants and in HopT1-1 and HopT1-1m3 transgenic plants relative to the miRNA accumulation in WT plants is shown for three biological replicates**. (E)** Flag-AGO1 was immunopurified from the inflorescence of the plants described in (D) and accumulation level of Flag-AGO1 proteins was assessed by western blot (top panel). The accumulation level of miRNAs associated to Flag-AGO1 in each condition was analyzed by Northern blot. Bottom panel: quantification of the miRNA accumulation level in HopT1-1 and HopT1-1m3 plants relative to the miRNA accumulation in *AGO1*_pro_::Flag-AGO1 transgenic plants is shown for three biological replicates

### HopT1-1 reduces the level of AGO1-bound miRNAs in a GW-dependent manner

The fact that HopT1-1 does not interfere with the biogenesis nor the stability of miRNAs (Fig. 6C), prompted us to assess whether it could, instead, inhibit the level of AGO1-bound miRNAs. To test this hypothesis, we transformed the previously described *AGO1*_pro_::FLAG-AGO1/*ago1*-*36* line (FLAG-AGO1) (Baumberger *et al*., 2005) with either the *35S*_pro_::HopT1-1 or *35S*_pro_::HopT1-1m3 constructs. Because AGO1 is well expressed in reproductive organs, we collected inflorescences from the above primary transformants expressing similar level of *HopT1-1* (Supplemental Fig. 6A), further immunoprecipitated AGO1, and assessed the levels of canonical miRNAs by low molecular weight northern blot analysis. Although we did not detect changes in the accumulation of miR156, miR159, miR160, miR168 and miR393 in the input samples, we consistently found a ∼2-fold decrease in the levels of immunoprecipitated AGO1-associated miRNAs from HopT1-1 expressing inflorescences compared to FLAG-AGO1 control inflorescences in three independent experiments (Fig. 6, D and E, Supplemental Fig. 6, B and C). By contrast, a similar level of these immunoprecipitated AGO1-bound miRNAs was recovered from HopT1-1m3 expressing inflorescences compared to FLAG-AGO1 inflorescences (Fig. 6, D and E, Supplemental Fig. 6, B and C). Altogether, these data provide evidence that HopT1-1 reduces the levels of AGO1-associated miRNAs in a GW-dependent manner, a molecular effect that likely contributes to the suppression of miRNA function triggered by this effector.

### HopT1-1 suppresses PTI responses in a GW-dependent manner and its presence mimics the impaired PTI responses observed in *ago1* mutants

Given that the majority of *Pto* DC3000 effectors promotes pathogenicity by dampening plant immune responses (Block and Alfano, 2011), we further investigated the ability of HopT1-1 to suppress PTI. Furthermore, the fact that HopT1-1 promotes pathogenicity through the genetic targeting of AGO1 (Fig. 1A, Supplemental Fig. 1, A and B) prompted us to assess the functional relevance of its AGO-binding platform in this process. To address these questions, we decided to use the EtHAn system, a recombinant *P. fluorescens* strain that triggers classical PTI responses and that expresses a functional type III secretion system, allowing the delivery of individual bacterial effectors in host cells (Thomas *et al*., 2009). It is noteworthy that this experimental system has been successfully used to characterize the PTI suppression activities of individual *Pto* DC3000 effectors during infection (Thomas *et al*., 2009). WT plants were infiltrated with EtHAn alone or with EtHAn strains expressing either HopT1-1 (EtHAn (HopT1-1)) or HopT1-1m3 (EtHAn (HopT1-1m3)), and the production of the reactive oxygen intermediate hydrogen peroxide (H_2_O_2_) was monitored at 24 hours post-inoculation by DAB staining (Fig. 7A). While the EtHAn strain induced strong production of H_2_O_2_ in WT plants, particularly within and around leaf vasculature, this phenotype was significantly reduced upon delivery of HopT1-1 (Fig. 7A). This result indicates that HopT1-1 can inhibit PAMP-triggered production of H_2_O_2_ in a physiological context of bacterial infection. By contrast, this PTI suppression effect was almost fully abolished upon delivery of the HopT1-1m3 mutant version (Fig. 7A), supporting a critical role for the GW motifs of HopT1-1 in this process. In addition, we exploited the EtHAn system to monitor the impact that HopT1-1, and its HopT1-1m3 mutant derivative, could have on bacterial-triggered deposition of callose, a late PTI response that plays a critical role in the establishment of basal immunity (Hauck *et al*., 2003). Interestingly, we found a ∼ 45% decrease in the number of callose deposits in response to EtHAn (HopT1-1) as compared to the EtHAn control, while this PTI suppression effect was not observed upon inoculation of the EtHAn (HopT1-1m3) strain (Fig. 7B). Taken together, these data indicate that HopT1-1 can suppress two well-established PTI responses during bacterial infection. They also provide evidence that the RNA silencing suppression activity of HopT1-1 is directly coupled with its ability to dampen PTI in Arabidopsis.

**Fig. 7.**
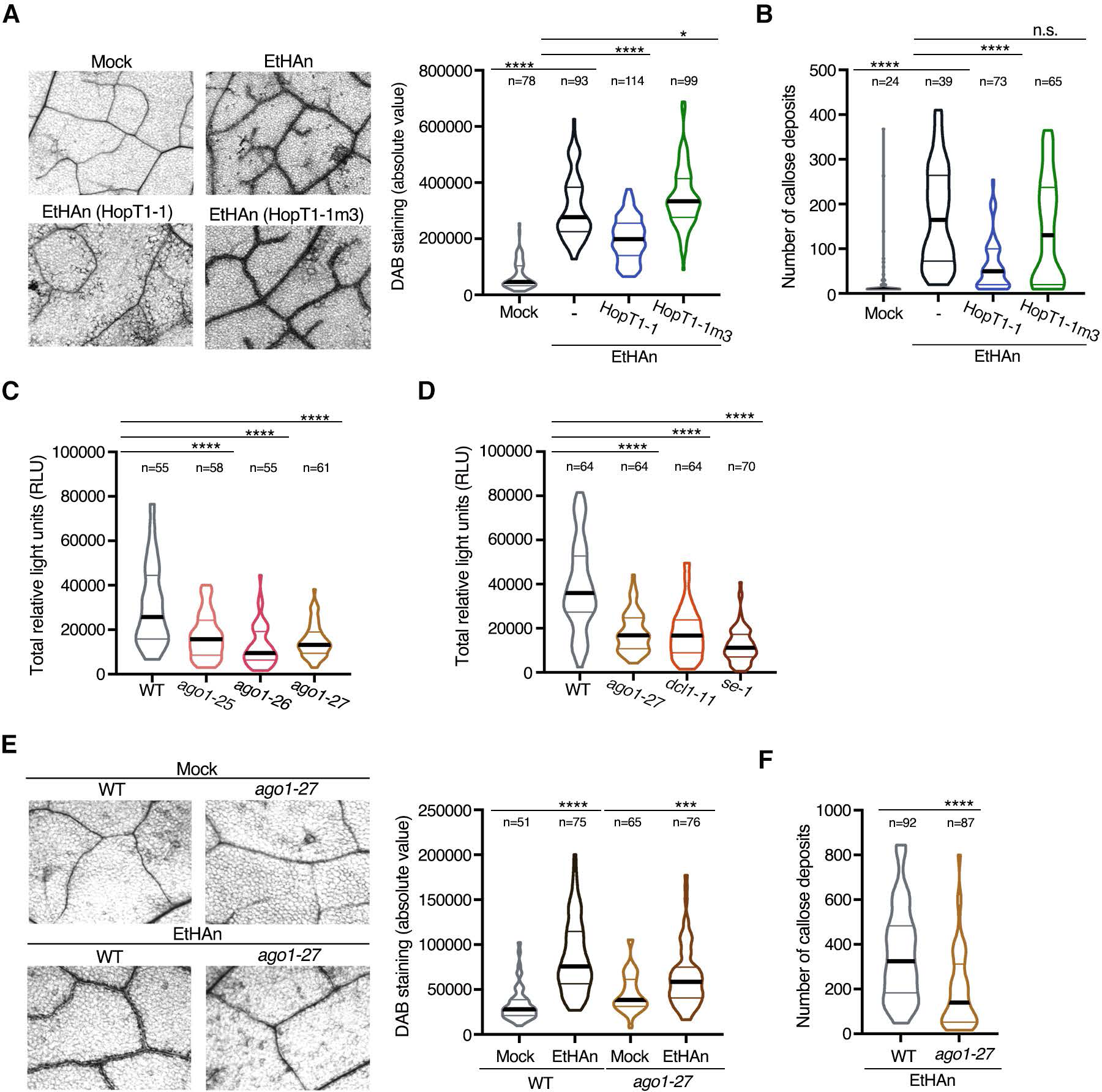
HopT1-1 dampens PTI in a GW-dependent manner and its presence mimics the impaired PTI responses observed in *ago1* mutants. **(A)** Detection of H_2_O_2_ production in the leaves of WT plants 24 hours after infiltration with the EtHAn strain alone (EtHAn) or with EtHAn strains carrying a plasmid encoding HopT1-1 or HopT1-1m3, respectively (left panel). Leaves from three plants were treated with ethanol to clear the chlorophyll pigments and were further incubated with DAB staining buffer to detect the presence of H_2_O_2_. Around 20-30 pictures were taken for each condition and absolute value of DAB staining was quantified using ImageJ software and presented as a violin plot (right panel). **(B)** Presence of callose deposits was detected 7 hours after infiltration of WT plants with the EtHAn strains used in (A). For each condition, leaves from three plants were collected and stained with aniline blue to detect the presence of callose deposits. The amount of callose deposits was measured using ImageJ software and presented as a violin plot. **(C)** Flg22-induced ROS production assay was performed in leaf discs from WT and *ago1* mutant alleles, *ago1-25, ago1-26 and ago1-27*. For each condition, leaves from three five-week-old plants were used to prepare leaf discs. Luminescence (Relative Light Unit; RLU) was subsequently measured for 45 min after flg22-elicitation for each technical replicate. The total amount of RLU produced during the flg22-elicitated time-course is presented as a violin plot with the mean as a horizontal bar. **(D)** As in (C) but in WT, *ago1-27*, *dcl1-11* and *se-1* mutants. **(E)** Same analysis as in (A) but in WT *versus ago1-27* mutant plants infiltrated with Mock or EtHAn. **(F)** Same analysis as in (B) but in EtHAninfiltrated WT plants versus *ago1-27* mutant plants. For all the above experiments, except for (E) and (F), statistical significance was assessed using one-way ANOVA analysis (*: p-value < 0.05; ****: p-value < 0.0001). For (E) and (F) experiments, statistical significance was assessed using Student’s *t* test (**: p-value < 0.01; ***: p-value < 0.001; ****: p-value < 0.0001). All the results shown in the different panels of the Fig. were pooled from at least two or three independent experiments. The number of samples (n) is indicated above each condition.

Given that AGO1 is a critical target of HopT1-1 (Fig. 1 and 3), the above findings suggested a major role for AGO1 in orchestrating PTI. To test this possibility, we first monitored flg22-triggered oxidative burst in the *ago1-25*, *ago1-26* and *ago1-27* hypomorphic mutant alleles. Importantly, all these *ago1* mutants displayed a compromised flg22-induced ROS production as compared to WT-elicited plants (Fig. 7C). A significantly impaired flg22-induced ROS production was also detected in the miRNA biogenesis defective mutants *dcl1-11* and *se-1* as compared to WT plants (Fig. 7D), indicating that the Arabidopsis miRNA pathway positively regulates this early PTI response. Hydrogen peroxide production and callose deposition were also reduced in *ago1-27* mutants *versus* WT plants challenged with the EtHAn strain (Fig. 7, E and F), although a milder effect was observed in this mutant background as compared to the effect detected in WT leaves challenged with the EtHAn (HopT1-1) strain (Fig. 7, A and B). These results support a role for AGO1 in PAMP-induced callose deposition, as previously reported during flg22 elicitation (Li *et al*., 2010). In addition, they provide evidence that AGO1 plays a central role in the production of ROS during bacterial elicitation. Altogether, these data suggest that the ability of HopT1-1 to suppress both PAMP-triggered ROS production and callose deposition involves, at least in part, an inhibitory effect of AGO1 activity.

## DISCUSSION

In this study, we have demonstrated a key role for the Arabidopsis miRNA pathway in PAMP-induced ROS production, a well-characterized PTI response that occurs within minutes of PAMP detection and that plays a crucial role in antibacterial defense (Couto and Zipfel, 2016; Torres *et al*., 2006). We have shown that the Arabidopsis miRNA factors DCL1, SE and AGO1 are all required for the first flg22-induced oxidative burst (Fig. 7). Furthermore, we found that the EtHAn strain triggers an intense accumulation of H_2_O_2_ within and around Arabidopsis leaf vasculature, a phenotype that was found partially impaired in the hypomorphic *ago1-27* mutant (Fig. 7). We propose that such AGO1-dependent regulatory process might ensure the formation of an immune cell layer adjacent to the vasculature to limit bacterial spreading from xylem vessels to mesophyll cells and *vice versa*. In addition, this phenomenon might result in an enhanced accumulation of H_2_O_2_ in xylem vessels, thereby potentially reducing bacterial survival through the well-characterized bactericidal function of this reactive oxygen intermediate (Hong *et al*., 2013). Such an antibacterial activity would notably be relevant to control *Pto* DC3000 pathogenesis, because this bacterium was previously shown to propagate through xylem vessels in both Arabidopsis and *N. benthamiana* (Yu *et al*., 2013; Misas-Villamil *et al*., 2011; Halter *et al*., 2021). Although the detailed mechanisms by which the Arabidopsis miRNA pathway orchestrates ROS production, and more generally PTI, remain to be established, the previously characterized miRNA-dependent control of negative regulators of PTI such as Auxin Response Factors 16 and 17 must contribute to this process (Li *et al*., 2010; Fig. 5 and 8). It is also equally possible that the previously described role of AGO1 in the transcriptional activation of hormone- and stress-responsive genes, including flg22-regulated genes, would contribute to this process (Liu *et al*., 2018). Furthermore, because the first PAMP-induced oxidative burst is an immediate immune signaling response, it is possible that the miRNA-directed control of ROS production might additionally be caused by a pool of pre-loaded AGO1-miRISC that would operate at the surface immune receptor complex, an intriguing possibility that will deserve attention in future studies.

**Fig. 8.**
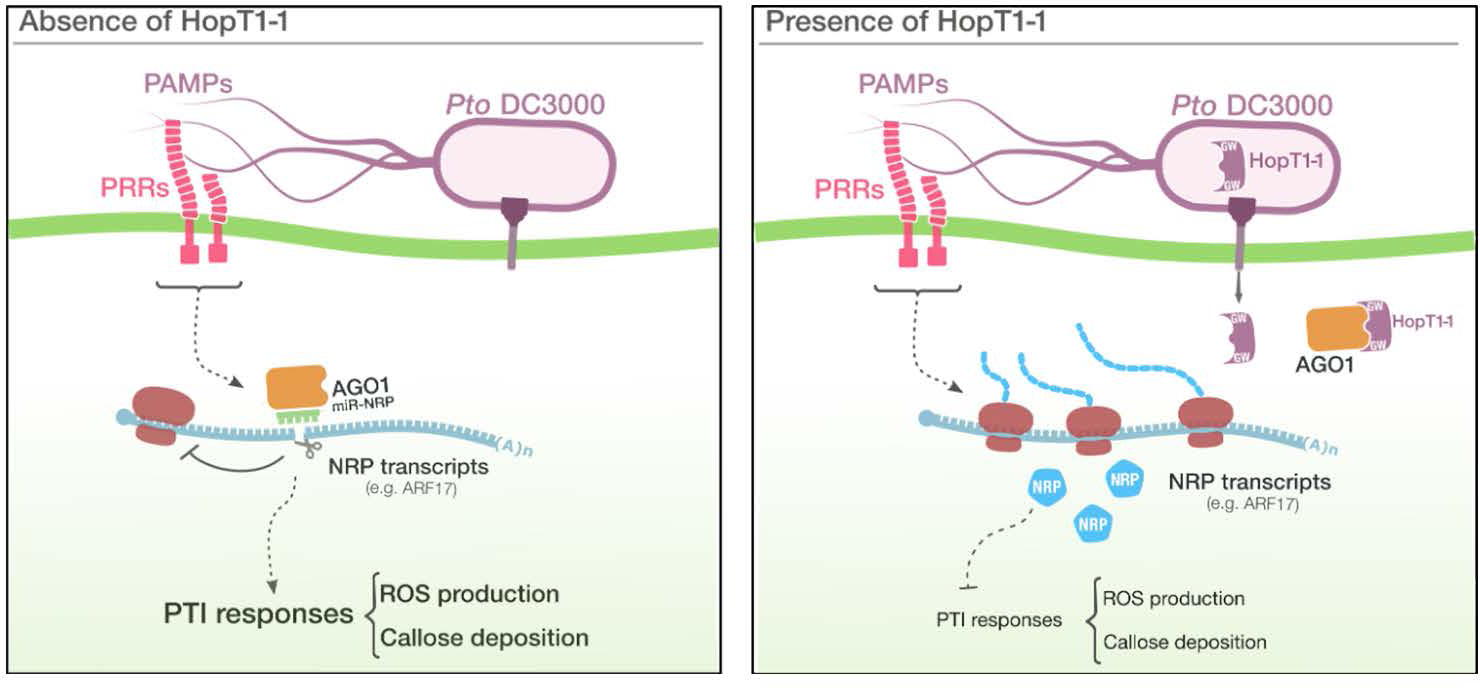
Hypothetical model for HopT1-1-triggered suppression of PTI. In the absence of HopT1-1: some functional Negative Regulator of PTI (NRPs) are silenced by miRNAs loaded in AGO1, a phenomenon that can be reinforced by the enhanced production of some miRNAs during PTI. This is notably the case of the NRP Auxin Response Factor 17 (ARF17) that is down-regulated during PTI through the upregulation of miR160a (Li *et al*. 2010). This regulatory mechanism is necessary to mount proper PTI responses, including PAMP-triggered ROS production and callose deposition. In the presence of HopT1-1: the injected *Pto* DC3000 effector HopT1-1 physically interacts with the two potentially functional W-binding pockets of the PIWI domain of Arabidopsis AGO1 through two conserved GW motifs. This phenomenon further alters the association and/or stability of miRNAs in the AGO1-miRISC, thereby derepressing some functionally relevant NRPs (*e.g*. ARF17). The enhanced accumulation of these NRPs further dampens PTI responses, including PAMP-triggered ROS production and callose deposition.

Given that the Arabidopsis miRNA pathway is a major component of PTI (Navarro *et al*., 2008; Li *et al*., 2010; this study), it is likely that many pathogen effectors will be found to target this small RNA pathway to enable disease. Consistent with this idea, we have previously identified type III secreted proteins from *Pto* DC3000 that suppress all the steps of the Arabidopsis miRNA pathway (Navarro *et al*., 2008). However, until now, it was unknown whether some of these BSRs could directly interfere with components of the RNA silencing machinery to suppress miRNA activity and cause disease. In the present work, we show that the bacterial effector HopT1-1 is a critical virulence determinant that promotes growth of *Pto* DC3000 in a physiological context of infection (Fig. 1). Importantly, the reduced growth of the *hopT1-1*-deleted strain was found fully rescued in miRNA-defective mutants including the hypomorphic *ago1-25, ago1-26* and *ago1-27* mutant alleles (Fig. 1), indicating that the Arabidopsis AGO1-dependent miRNA pathway is a major genetic target of HopT1-1 during infection. In agreement with these functional analyses, we found that HopT1-1 can physically interact with Arabidopsis AGO1 through two GW motifs (Fig. 8), which were found conserved in HopT1-1 homologs from various bacteria, including the distant bacterium *Marinomonas mediterranea* (Fig. 2). Using structural modeling and manual docking analyses, we further showed that the W112 motif of HopT1-1 unveiled sensible assemblies with the two potentially functional W-binding pockets of the PIWI domain of Arabidopsis AGO1 (Fig. 4). Interestingly, we also noticed that the HopT1-1 molecules interacting with these pockets occupy different regions of the three-dimensional space around Arabidopsis AGO1 (Fig. 4). The Arabidopsis PIWI domain of AGO1 can therefore potentially accommodate two HopT1-1 molecules simultaneously, one in each W-binding pocket (Fig. 4). In addition, we showed that the AGO-binding platform of HopT1-1 plays a central role in the suppression of miRNA activities, as reflected by the derepression of unrelated endogenous miRNA targets, which was specifically observed in HopT1-1 but not in HopT1-1m3 primary transformants (Fig. 5). Such AGO-binding platforms were also found essential for the suppression of PAMP-triggered H_2_O_2_ production and of callose deposition in Arabidopsis (Fig. 7 and 8). Altogether, these data imply that the BSR activity of HopT1-1 is directly coupled with its ability to suppress PTI. This phenomenon might be due to HopT1-1-induced derepression of known negative regulators of PTI that are controlled by miRNAs (Fig. 8; Li *et al*., 2010). It could also be due to HopT1-1-triggered suppression of (i) the transcriptional reprogramming activity of the chromatin-bound pool of AGO1 during PTI (Liu *et al*., 2018), and/or (ii) the above hypothesized AGO1-directed early PTI signaling events at the surface immune receptor complex.

To understand the mechanisms by which HopT1-1 suppresses the Arabidopsis miRNA pathway, we further profiled total small RNAs in HopT1-1 and HopT1-1m3 transgenic plants. Importantly, we did not detect a general decrease in the accumulation of mature miRNAs in HopT1-1 transgenic plants (Fig. 6, A and B). Accordingly, the levels of canonical miRNAs, tested by northern blot analysis, did not show significant changes in HopT1-1 transgenic plants compared to WT or HopT1-1m3 transgenic plants (Fig. 6, C and D). This was also true for a subset of primary miRNA transcripts, whose levels were either unchanged or barely changed in HopT1-1 transgenic plants compared to WT plants and HopT1-1m3 transgenic plants (Supplemental Fig. 5). On the contrary, we found that HopT1-1 consistently reduced the level of canonical miRNAs in AGO1 immunoprecipitates, a phenomenon which was found to be dependent on its AGO-binding platform (Fig. 6E). Based on these data, we propose that HopT1-1 must act at the level of the AGO1-miRISC, rather than at the levels of miRNA biogenesis or stability (Fig. 8). This work therefore suggests that the silencing suppression activity of HopT1-1 over Arabidopsis miRNA targets is caused by its ability to bind AGO1 through its GW motifs, and to further alter the association of miRNAs with AGO1 and/or their stability in this silencing effector complex. Future investigations will be required beyond this study to determine the detailed underlying mechanisms. It will notably be appealing to determine whether HopT1-1 could prevent and/or displace the interaction of AGO1 with yet-unknown Arabidopsis GW/WG co-factor(s) of AGO1, that would regulate the loading and/or stability of miRNAs at the AGO1-miRISC level.

The present study reveals that the use of GW/WG-dependent AGO-binding platforms is not restricted to VSRs (El-Shami *et al*., 2007; Azevedo *et al*., 2010; Garcia *et al*., 2012; Karran and Sansfaçon, 2014; Aqil *et al*., 2013; Giner *et al*., 2010), but can also be exploited by a functionally relevant bacterial effector. Intriguingly, we have also retrieved GW/WG motifs in an appropriate sequence context in secreted effectors from various phytopathogenic bacteria, fungi and oomycetes (Supplemental Fig. 7). For instance, we have identified the presence of canonical GW/WG motifs in effectors from the devastating plant bacterial vascular pathogens *Xylella fastidiosa*, *Xanthomonas campestris* pv. *campestris* and *Xanthomonas oryzae pv. oryzae,* or from the oomycetes *Phytophthora sojae* and the Irish potato famine pathogen *Phytophthora infestans* (Supplemental Fig. 7). Canonical GW/WG motifs were also retrieved in effectors produced by the wheat stem rust pathogen *Puccinia graminis* f. sp. *tritici*, which represents one of the most destructive fungal pathogen of wheat worldwide, and that was shown to encode functional fungal suppressors of RNA silencing (FSRs) (Yin *et al.,* 2019), but also by *Fusarium graminearum*, which is the causal agent of Fusarium head blight that causes serious yield losses in wheat and barley worldwide (Supplemental Fig. 7). Collectively, these observations suggest that a wide range of non-viral phytopathogens might have evolved an analogous mode of action to promote pathogenicity in agriculturally relevant crops. It will thus be interesting to establish if the discoveries made on HopT1-1 hold for other silencing suppressors from non-viral phytopathogens. It will also be worth developing innovative strategies to counteract this potentially widespread pathogen-mediated virulence mechanism in order to confer broad spectrum disease resistance in crops.

## MATERIALS AND METHODS

### DNA Constructs

The pK7WG2D destination vector carrying the *HopT1-1* gene has been previously described (Navarro *et al*., 2008). To generate the mutated versions of *HopT1-1*, GW>GF substitutions in the *HopT1-1* ORF were carried out by PCR-based site-directed mutagenesis using mismatched primers. The resulting PCR product was introduced in the pENTR/D-TOPO entry vector (Invitrogen, Carlsbad, CA), sequenced and then recombined into the GATEWAY binary destination vector pK7WG2D using LR clonase (Invitrogen, Carlsbad, CA), to overexpress an untagged protein, or into the GATEWAY binary destination vector pEarleyGate 203 to overexpress a Myc fusion protein. Both of the plasmids allow HopT1-1 expression under the control of the constitutive CaMV 35S promoter (*35S*_pro_). To overexpress HopT1-1 and HopT1-1m3 in the EtHAn system (Thomas *et al*., 2009), versions of *HopT1-1* and *HopT1-1m3* without a stop codon were cloned in the pENTR/D-TOPO entry vector, which was further recombined with the destination vector pBS0046 containing the *NPTII* promoter. For the BiFC assays, the coding regions of *AGO1*, *SDE3*, *HopT1-1*, *HopT1-1m3* and *HopC1* were first cloned into the pDONR207 plasmid and further recombined into the GATEWAY BiFC binary destination vectors pUBN-YFP, containing the N-ter or the C-ter part of YFP protein, that were previously described in Grefen *et al*. (2010). For the FRET-FLIM experiment, the no stop versions of pDON-HopT1-1 and pDON-HopT1-1m3 were recombined into the destination vector pBin-35S-GWY-YFP. For Agrobacterium-mediated transformation in *N. benthamiana*, pDON-HopT1-1 and pDON-HopT1-1m3 were recombined with the GATEWAY pEarleyGate 203 plasmid expressing a MYC tag in N-ter. All the constructs inserted in GATEWAY binary destination vectors were transformed into the *Agrobacterium tumefaciens* strains GV3101 or C58C1.

### Transgenic plants and mutants

*Arabidopsis thaliana* silencing defective mutants *ago1-25, ago1-26, ago1-27*, *ago2-1*, *ago4-2*, *ago4-3*, *dcl2-1 dcl4-2*, *se-1*, *dcl1-11*, *sgs3-1*, *rdr1-1 rdr2-1 rdr6-15* were previously described (Morel *et al*., 2002; Agorio and Vera, 2007; Havecker *et al*., 2010; Henderson *et al*., 2006; Zhang *et al*., 2008; Mourrain *et al*., 2000; Garcia-Ruiz, 2010; de Felippes *et al*., 2011). To generate plants constitutively expressing HopT1-1 and HopT1-1m3 under the control of the 35S promoter, pK7WG2D vectors carrying *HopT1-1* or *HopT1-1m3* sequences were transformed in Col-0 (WT plants) using Agrobacterium-mediated transformation and multiple individual T1 primary transformants (12 or 13 individuals) were analyzed in parallel of WT individual plants.

### Bacterial strains

The bacterial strains used in this study include *Pto* DC3000, *HopT1-1*-deleted strain (*Pto* Δ*HopT1-1*) and *HopC1*-deleted strain (*Pto ΔHopC1*). These bacterial strains were grown at 28°C in NYGB medium (5 g L^-1^ bactopeptone, 3 g L^-1^ yeast extract, 20 ml L^-1^ glycerol) containing rifampicin (25 µg/mL) for selection. The *Pto ΔHopT1-1* was generated by an unmarked mutagenesis strategy described by House *et al*. (2004) with some modifications. Briefly, DNA regions upstream and downstream of *HopT1-1* were amplified with primer pairs P4835/P4836 and P4837/P4838 (Table S1), respectively. The resulting PCR fragments were cloned into pENTR/D-TOPO and then recombined into suicide destination vector pMK2016 for the upstream construct, or pMK2017 for the downstream construct, using LR Clonase according to the manufacturer’s instructions, resulting in constructs pLN5426 and pLN5472, respectively. Constructs pLN5426 and pLN5472 were integrated into the *Pto* DC3000 chromosome by biparental or triparental mating. Plasmid pBH474, which contains a gene encoding the yeast FLP recombinase, was introduced into the resulting strains to excise the integrated plasmid at FLP recombinase target (FRT) sequences. To cure pBH474, isolated colonies were spread on King’s B medium (King *et al.,* 1954) agar plates containing rifampicin (100 µg/mL) with 5% sucrose. PCR primer pair P4839/P4840 (Table S1) was used to identify mutants that carried the *HopT1-1* deletion. The *Pto ΔHopC1* was generated by an insertional mutagenesis strategy described by Windgassen *et al*. (2000) with some modifications. Briefly, an internal fragment within the *HopC1* coding region was amplified with primer pair P164/P165 (Table S1). The resulting PCR fragment was cloned into a suicide vector pKnockout-Ω that had been digested with XcmI, resulting in construct pLN6. Construct pLN6 was integrated into the *Pto* DC3000 chromosome by triparental mating using spectinomycin (50 μg/mL) to select for the plasmid marker. PCR primer pair P181/P165 was used to identify mutants that contained an integrated pKnockout-Ω in *HopC1*.

### Plant Growth Conditions and Treatments

Plants used in this study were grown in growth cabinets at 19–23 °C (dark/light) with an 8-h photoperiod for all the assays. Transgenic plants expressing HopT1-1, under the control of the constitutive CaMV 35S, were selected on plates containing Murashige and Skoog medium (Duchefa) [composition for a 1-L medium (pH = 5.7): 2.3 g MS, 0.5% sucrose, 0.8% agar, 0.5 g MES, vitamins (Sigma)] in presence of Kanamycin (50 μg/mL) and then transferred to soil 15 day-post-germination (dpg). *Nicotiana benthamiana* plants for transient expression assays were grown in growth chamber at 19-23°C (dark/light) with a 16-h photoperiod.

### Bacterial growth assays

Five-week-old Col-0 Arabidopsis (WT) or the mutant plants were covered with a lid for 24 hours under high humidity before being dip-inoculated with *Pto* DC3000 WT, *Pto ΔHopT1-1* or *Pto ΔHopC1* at 10^8^ cfu/mL with 0,02% of Silwet L-77 (Lehle Seeds). Dip-inoculated plants were allowed to stay under high humidity condition for 3 days before the bacterial counting was performed. For each condition, around 15 leaves from three plants were collected and the following steps were performed as described in Navarro *et al*. (2008).

### Agrobacterium-mediated transient expression

*Agrobacterium tumefaciens* strains carrying the indicated plasmids were inoculated in 10 mL of LB medium supplied with rifampicin and the other selective antibiotic, and placed in an incubator shaker (200 rpm) at 28°C overnight. Cells were harvested by centrifugation at 4500 rpm and resuspended to a final optical density at 600 nm (OD_600_) of 0.2 in a solution containing 10 mM MES, pH 5.6, 10 mM MgCl_2_ and 200 μM acetosyringone. Cultures were incubated under agitation in the dark at room temperature for 5 hours before agro-infiltration. For the co-infiltration of two different *A. tumefaciens* strains, equal concentrations (OD_600_: 0.25) of both cultures were mixed before agroinfiltration of leaves of four-week-old *Nicotiana benthamiana* plants. Infiltrated plants were covered with a lid and the leaves were collected and flash frozen 48-72 hours post-infiltration. Confocal image analyses for the BiFC experiments were performed 2-3 days post infiltration. YFP fluorescence for each sample was visualized and acquired by using a Leica SP8 confocal microscope with a 63x oil immersion objective, and quantification of the fluorescence signal for each picture taken (n=10) was performed using ImageJ software (National Institutes of Health, Bethesda).

### Quantitative Real Time PCR analysis

Total RNA was extracted using a phenol-based extraction protocol (Box *et al*., 2011) followed by DNAse (Promega) digestion at 37°C to remove the genomic DNA. 0.5 µg of DNAse-digested RNA were reverse-transcribed into cDNA using qScript cDNA Supermix (Quanta Biosciences). The cDNA was quantified using a SYBR Green qPCR mix (Takyon; Eurogentec) and gene-specific primers. PCR was performed in 384-well plates heated at 95 °C for 10 min, followed by 45 cycles of denaturation at 95 °C for 10 s and annealing at 60 °C for 40 s. A melting curve was performed at the end of the amplification. Transcript levels were normalized to that of *Actin2* and/or *Ubiquitin5* levels. Primers used to monitor *Actin2*, *SPL10*, *ARF17, PHB, DCL1, ARF10, pri-miR159b, pri-miR166a, pri-miR167a* and *Ub5* were previously described (Navarro *et al*., 2008; Zhang *et al*., 2008; Vaucheret *et al.,* 2004; Vazquez *et al*., 2004; Cai *et al.,* 2018; La Camera *et al*., 2009). Primer pairs used for the remaining genes are listed in Table S1.

### RNA gel blot analyses

Total RNA was extracted from Arabidopsis tissues (5-week-old plants) with Tri-Reagent (Sigma, St. Louis, MO) according to the manufacturer’s instructions. RNA gel blot analysis of low molecular weight RNAs was performed on 15-20 μg of total RNAs and as described previously (Navarro *et al*., 2008). Detection of U6 RNA was used to confirm equal loading. DNA oligonucleotides complementary to miRNA sequences were end-labeled with γ-^32^P-ATP using T4 PNK (New England Biolabs, Beverly, MA).

### Protein extraction and analyses

Total protein extracts were done according to Azevedo *et al*. (2010) and resolved on SDS-PAGE after quantification with Bradford Assay. After electroblotting proteins on nitrocellulose membrane (Millipore), protein blot analysis was performed using antiserum with specific antibodies. Antibodies against AGO1 (AS09527) and PEPC (AS09458) were purchased from Agrisera.

### Pull-down experiments

Biotinylated peptides (synthetized by Eurogentec, sequences shown in Fig. 2) were mostly insoluble in water and were thus resuspended in 6 M urea, and sonicated four times (Bioruptor^TM^, Diagenode, 30s on/1min off on High). They were quantified at 205 nm using the Nanodrop 2000 according to the manufacturer’s instruction (Thermoscientific) and checked by dot blot analysis. The solubilized peptides were spotted on a nitrocellulose membrane at three different amounts (1 µg, 0.1 µg and 0.01 µg) and were detected by using the streptavidin protein conjugated to HRP (21126, Thermoscientific), and the ECL substrate. For binding assays, 10 µg of peptides were diluted into 450 µL of PBS containing 0.1% of NP-40 and were incubated for 30 min at room temperature, in the presence of 0.9 mg of Dynabeads® MyOne™ Streptavidin T1. The beads were then washed once in 0.1% NP-40 PBS and twice in IP buffer (10% glycerol, 50 mM Tris-HCl pH 7.4, 150 mM NaCl, 5 mM MgCl_2_, 0.1% NP-40, 2 mM DTT, EDTA-free complete Protease Inhibitor Cocktail (Roche)). They were incubated in the presence of inflorescence extracts from Flag-AGO1 Arabidopsis plants in IP buffer for 2 hours at 4°C. After 3 washes of the beads in IP buffer, the proteins were eluted in Laemmli buffer and resolved on a 6% SDS-PAGE gel. The AGO1 protein was detected by Western blot using a Flag-HRP antibody (A8592, Sigma).

### FRET-FLIM Measurement

Fluorescence lifetime measurements were performed in time domain using a streak camera (Camborde *et al.,* 2017). The light source is a 440 nm pulsed laser diode (PLP-10, Hamamatsu, Japan) delivering ultrafast picosecond pulses of light at a fundamental frequency of 2 MHz. All images were acquired with a 60x oil immersion lens (plan APO 1.4 N.A., IR) mounted on an inverted microscope (Eclipse TE2000E, Nikon, Japan). The fluorescence emission is directed back into the detection unit through a short pass filter and a band pass filter (490/30 nm). The detector is a streak camera (Streakscope C4334, Hamamatsu Photonics, Japan) coupled to a fast and high-sensitivity CCD camera (model C8800-53C, Hamamatsu). For each acquisition, average fluorescence decay profiles were plotted and lifetimes were estimated by fitting data with exponential function using a non-linear least-squares estimation procedure (Camborde *et al*., 2017). Fluorescence lifetime of the donor was experimentally measured in the presence and absence of the acceptor. FRET efficiency (E) was calculated by comparing the lifetime of the donor in the presence (ρ_DA_) or absence (ρ_D_) of the acceptor: E=1- (ρ_DA_)/(ρ_D_). Statistical comparisons between control (donor) and assay (donor + acceptor) lifetime values were performed by Student’s *t* test.

### Quantification of flg22-induced ROS production

For each condition, discs (0.4 cm in diameter) of leaf tissue were harvested from three individual five-week-old plants and incubated in water in 96-well plates overnight in a growth chamber at 23°C. After 24 hours, the water was removed and replaced by 100 µL of H_2_O containing 20 µM luminol and 1 µg of horseradish peroxidase (Sigma) with 100 nM flg22. Luminescence (relative light units) was immediately measured for a timecourse of 45 min using a Tristar LB 941 plate reader (Berthold technologies).

### Callose deposition

Five-week-old Arabidopsis Col-0 (WT) plants or *ago1-27* mutant plants were infiltrated with 10 mM of MgCl_2_ or 10^8^ cfu/mL (OD_600_: 0.2) of EtHAn strains. Seven hours after infiltration, around 12 leaves were collected from three independent plants and incubated overnight in lactophenol (1 volume of glycerol:lactic acid:phenol:water, ratio 1:1:1:1, and 2 volumes of EtOH). After a first washing step in 50% EtOH and a second one in water, leaves were incubated for 30 min in aniline blue staining solution (150 mM K_2_HPO_4_ pH 9.5 with 0.01% aniline blue). Leaves were mounted with 50% glycerol and visualized with Olympus Macro Zoom System Microscope MVX10 fluorescent microscope (excitation filter 365 nm and barrier filter 420 nm). The number of callose deposits was quantified using ImageJ software. Forty fields of view (each 0.56 mm^2^) were analyzed and averaged.

### DAB staining assay

Five-week-old Arabidopsis Col-0 (WT) plants or *ago1-27* mutant plants were infiltrated with 10 mM of MgCl_2_ or 10^8^ cfu/mL (OD_600_: 0.2) of EtHAn strains. After 48 hours, the infiltrated leaves were collected and vacuum-infiltrated with DAB staining buffer (1 mg/mL, pH 3.5) and then incubated for 5 hours in the same buffer. Leaves were boiled for 15 min in an EtOH:glycerol:acid lactic (ratio 4:1:1) solution, washed overnight in EtOH and then mounted with 50% glycerol and further visualized using Olympus Macro Zoom System Microscope MVX10. The intensity of DAB staining was quantified with ImageJ software. Forty fields of view (each 0.56 mm^2^) were analyzed and averaged.

### Trypan blue staining

Leaves from five 5 week-old Arabidopsis Col-0 (WT) plants or T2 transgenic lines expressing HopT1-1 or HopT1-1m3 were placed in a 15 mL tube and immersed in lactophenol trypan blue solution (acid lactic:glycerol:phenol:water:trypan blue, ratio 1:1:1:1; diluted before use at a ratio 1:1 with EtOH). The tube was placed into a boiling water bath for 2 min followed by a first incubation in 5 mL chloral hydrate solution (2.5 g/mL water) for 2 hours, and a second one overnight on an orbital shaker. Note that the staining step should not exceed the time indicated to avoid a strong background signal. After extensive washes in water, the leaves were mounted onto glass microscope slides in presence of 50% glycerol and examined using Olympus Macro Zoom System Microscope MVX10.

### Structural analysis of HopT1-1 and AGO1

For HopT1-1, the coordinates of AlphaFold models were obtained from the AlphaFold Protein Structure Database (Jumper *et al*., 2021; Varadi *et al*., 2022) (https://al-phafold.ebi.ac.uk/ with accession numbers Q88BP7, F2JYU5, A0A0P9KI05, A0A2A2E7U8, A0A3M5KIL0, Q87WF7). Structures were colored according to the AlphaFold per-residue confidence score and analyzed with PyMOL (https://www.pymol.org/). For Arabidopsis AGO1, the coordinates of the AlphaFold model were obtained from the AlphaFold Protein Structure Database (https://al-phafold.ebi.ac.uk/, accession number 004379). The Arabidopsis AGO1 AlphaFold model was superimposed on the tryptophane-bound structure of human Ago2 (pdb code 6cbd) (Sheu-Gruttadauria *et al*., 2018) using the align command in PyMOL, and tryptophane-binding pockets (Trp-pockets) of human Ago2 and Arabidopsis AGO1 were inspected. To manually dock HopT1-1 into Arabidopsis AGO1 putative Trp-pockets, AGO1 and HopT1-1 AlphaFold models were first prepared by removing segments that may be unstructured (pLDDT scores < 50). In Arabidopsis AGO1, this corresponds to segments 1-173, 276-281, and 999-1028, and in HopT1-1 to segments 1-27, and 89-108. In addition, a long segment of HopT1-1 (segment 185-252) that is not well-predicted in *Pseudomonas syringae* Pto DC3000 and that may be unstructured in some HopT1-1 members of the family (pLDDT scores < 50) was also removed. In Arabidopsis AGO1 model, the conformation of the side-chain of R865 was changed to that of R688 in human Ago2 in order to open the putative Trp-pocket-3 and make it available for interacting with GW motifs. The side chains of W112 and W181 were then manually aligned onto the tryptophane molecules of the 6cbd structures placed into the AtAGO1 Trp-pockets (Trp-pocket-1 and Trp-pocket-3) using the pair_fit command in PyMOL. Complexes without extensive structural overlaps between Arabidopsis AGO1 and HopT1-1 were further analyzed.

### Predicting the effect of GW mutations on HopT1 protein stability

In order to predict the effect of GW mutations on the stability of HopT1-1, the coordinates of AlphaFold models (https://alphafold.ebi.ac.uk/ with accession numbers Q88BP7, F2JYU5, A0A0P9KI05, A0A2A2E7U8, A0A3M5KIL0, Q87WF7) were provided to both PoPMuSiC and HoTMuSiC softwares (Dehouck *et al*., 2011; Pucci *et al*., 2016) as input. For each mutation, the average value as well as the standard deviation for the set of predicted values for identical mutations in the ensemble of AlphaFold models of the HopT1-1 family are reported. Mutations tested correspond to W79F, W79A, W112F, W112A, W181F, and W181A.

### Small RNA and RNA sequencing library analyses

RNAseq reads generated for this study were mapped to the Arabidopsis Col-0 genome (v TAIR10) with HopT1 and HopT1-1m3 sequences using STAR (v2.5.2b) (Dobin *et al*. 2013) with parameters --twopassMode Basic. Gene counts were obtained with HT-seq count (v 0.6) (Anders *et al*. 2015). Differential expression was assessed using DeSEQ2 (Love *et al*. 2014) with adjusted p-value <0.05. sRNA libraries were mapped against the *Arabidopsis thaliana* genome (v TAIR10) using ShortStack (Axtell, 2013) and Bowtie2 (Langmead and Salzberg, 2012) with parameters --mismatches 0 --dicermin 15 –dicermax 30 --bowtie_m 20 --mmap f. MicroRNA annotations were obtained from miRBase Release 22.1 (Kozomara *et al*. 2019). Graphics of the Fig6 A and B and Supplemental Fig. 4A were generated in R using packages ggplot2 (Wickham *et al*., 2011) and ggsignif (Gao *et al*., 2021). Raw reads from sRNA and RNA-seq libraries have been deposited to the ncbi SRA under BioProject “PRJNA995567”.

### Phylogeny and sequence analyses

Amino acid sequences of HopT1-1 orthologs (Group 7800626at2) were obtained from OrthoDB v11 (Kuznetsov *et al*. 2023). Multiple alignments were generated with Muscle (v3.7) (Edgar *et al*. 2004) and phylogenetic trees with PhyML (v 3.0) (Guindon *et al*. 2010) using the Phylogeny.fr platform (Dereeper *et al*. 2008). Alignements and trees were plotted in R with ggtree (Yu *et al*. 2017).

## Author contributions

O.T., L.N. designed research; O.T., M.S.R., M.C., F.Y., D.P., C.P., L.B., G.L. performed research; A.L.P-Q., D.A. performed bioinformatic analyses; P.B. performed structural modeling; O.T., M.S.R., L.D., T.L., J.R.A., A.L.P-Q., P.B., L.N. analyzed data; O.T., L.N. wrote the paper; T.L., D.P., L.D., C.P., J.R.A., L.N. acquired funding; all the authors reviewed and edited the paper.

## Acknowledgements

We thank J. Chang for the EtHAn strain, O. Voinnet for DCL1, AGO2 and AGO4 antibodies, P. Genschik for the CFP-AGO1 construct and all the members of the Navarro Lab for critical reading of the manuscript. We are grateful to the PlantAlgae Facility of IBENS, which received support from the program “Investissements d’Avenir” ANR-10-Labx-54 MEMOLIFE and ANR-11-IDEX-0001-02 PSL* Research University and of the SESAME Program from the “Région Île-de-France”; the Imaging Facility of IBENS, which received the support of grants from the “Région Île-de-France” (NERF N°2011-45), the “Fondation pour la Recherche Médicale” (N° DGE 20111123023) and the “Fédération pour la Recherche sur le Cerveau - Rotary International France” (2011), and the TRI-Genotoul platform, which was supported by the “Région Occitanie/Pyrénées-Méditerranée” (PRISM-Project). This work was supported by an European Research Council starting grant entitled “Silencing & Immunity” (to L.N.), an ATIP-Avenir Grant from the Fondation Bettencourt Schueller (to L.N.), a grant from the National Institutes of Health (to J.R.A.), and support from the Center for Plant Science Innovation at the University of Nebraska (to J.R.A.), by an ANR grant (08-BLAN-0206) and support from the CNRS (to T.L. and D.P.), by an ANR grant (15-CE20-0016-01) and support from the French Laboratory of Excellence ‘TULIP’ (ANR-10-LABX-41; ANR-11-IDEX-0002-02) (to L.D.) and by the “Région Occitanie/Pyrénées-Méditerranée” PRISM-Project (to C.P.).

