## Supplemental Figure 1 for "A bacterial effector directly targets Arabidopsis Argonaute 1 to suppress Pattern-triggered immunity and cause disease"

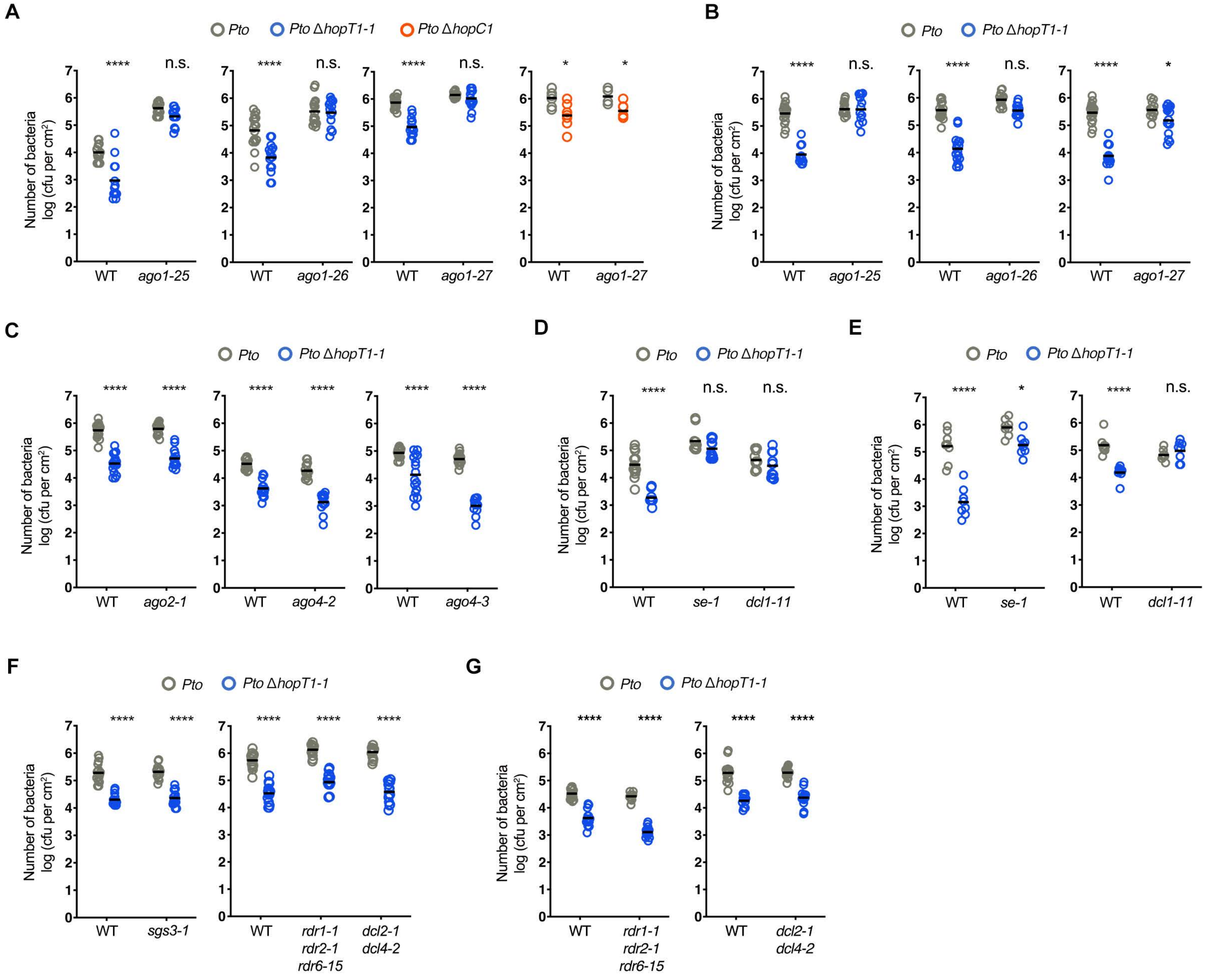


**Supplemental Figure 1. The growth defect of the *Pto* ΔhopT1-1 strain is specifically rescued in Arabidopsis miRNA-defective mutants**

Five-week-old Col-0 Arabidopsis (WT) plants and indicated genotypes in each panel were dip-inoculated with bacterial strain Pto DC3000 (Pto) (blue dot), Pto ΔhopT1-1 (green dot) or Pto ΔhopC1 (orange dot) at a concentration of 108 cfu/mL. At three days post-inoculation, leaves from three plants were collected and bacterial titers monitored. Each symbol represents a number of bacteria as log (cfu per cm2) and mean (n=8 or 16) is represented as a horizontal line in the dot plots. Independent biological replicates distinct from the one presented in Figure 1 are presented here. (A)-(B) Left panel: Growth of WT Pto DC3000 strain (Pto) and of hopT1-1-deleted bacterial strain (Pto ΔhopT1-1) in WT plants and in three different alleles of ago1 mutants, namely ago1-25, ago1-26 and ago1-27. Right panel: Growth of WT Pto DC3000 strain (Pto) and of hopC1-deleted bacterial strain (Pto ΔhopC1) in WT plants and in ago1-27 mutant. (C) Growth of WT Pto DC3000 and of Pto ΔhopT1-1 in WT plants and in mutant defective in AGO2 (ago2-1) or in AGO4 (ago4-2 and ago4-3). (D)-(E) Same as in (A-Left panel) but in miRNA biogenesis-defective mutants: se-1 and dcl1-11. (F)-(G) Same as in (C) but in siRNA biogenesis-defective mutants: sgs3-1, rdr1-1 rdr2-1 rdr6-15 and dcl2-1 dcl4-2. Statistical significance for all the above experiments was assessed using the ANOVA test (*: p-value < 0.05; **: p-value < 0.01; ***: p-value < 0.001; ****: p-value < 0.0001).
