## Supplemental Figure 2 for "A bacterial effector directly targets Arabidopsis Argonaute 1 to suppress Pattern-triggered immunity and cause disease"

A

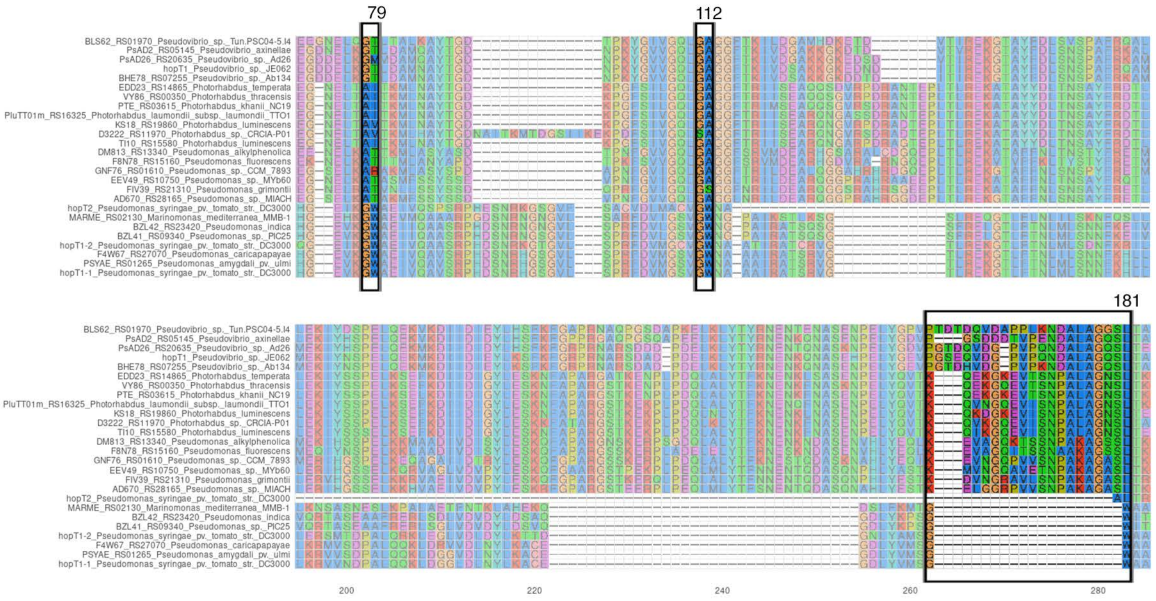


B

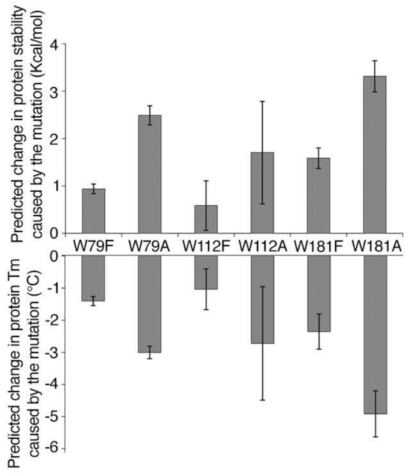


**Supplemental Figure 2. HopT1-1 possesses three conserved W residues that can be mutated in F residues with limited destabilizing effects**
**(A)** Complete sequences of the Pto DC3000 HopT1-1 family were aligned using Muscle. Each GW motifs is indicated by a black rectangle. **(B)** Prediction of changes in protein stability caused by protein mutations as reported by PoPMuSiC (v3.1) and HoTMuSiC (v1.0) software. Predicted changes in Gibbs free energy of folding (ΔΔG) in kcal/mol are reported in the upper part of the graph and predicted changes in protein melting temperature (ΔTm) are reported in the lower part. Positive ΔΔG and negative ΔTm correspond to destabilizing mutations. For each indicated mutated (W mutated to A or to F) residue, the average value as well as the standard deviation for the set of predicted values for identical mutations in the ensemble of AlphaFold models of the HopT1-1 family of panel B are reported. Tryptophane to phenylalanine mutations (i.e. W79F, W112F, and W181F) are predicted to have limited destabilizing effects but tryptophane to alanine mutations are predicted to lead to larger destabilizing effects.
