## Supplemental Figure 3 for "A bacterial effector directly targets Arabidopsis Argonaute 1 to suppress Pattern-triggered immunity and cause disease"

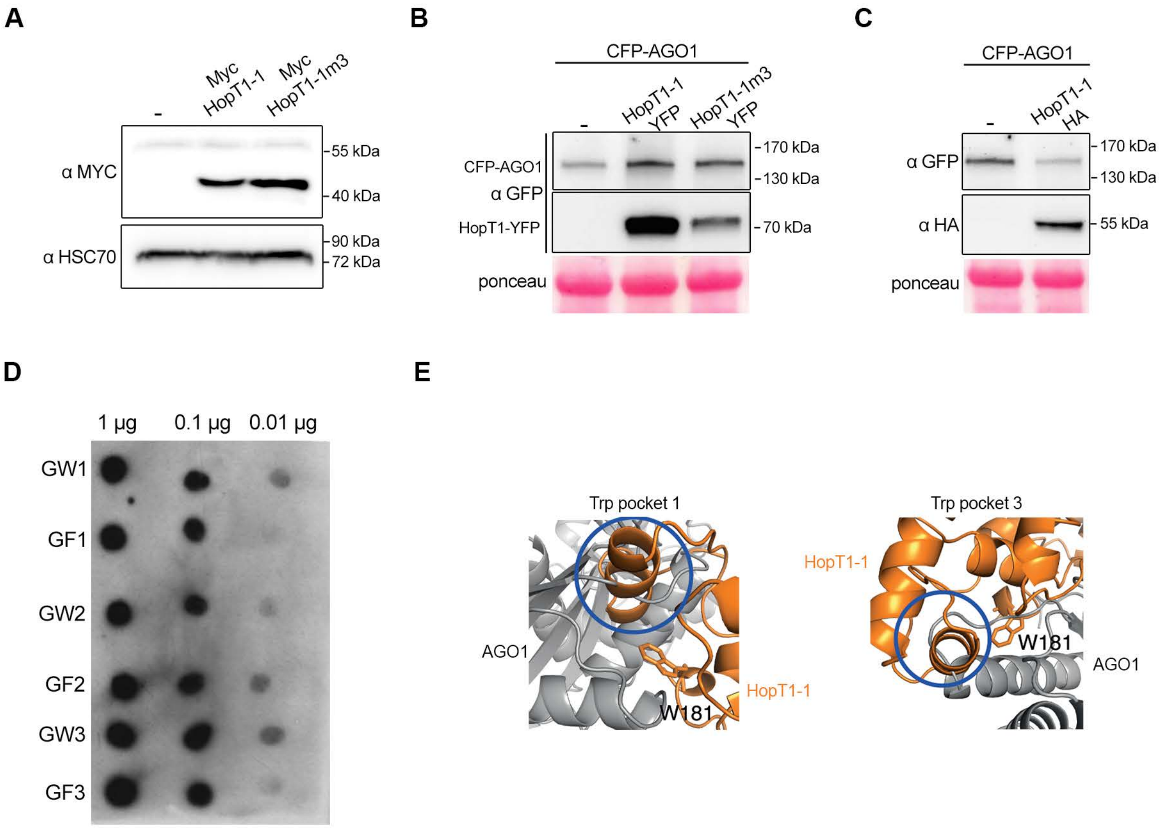


**Supplemental Figure 3. Accumulation level of full length HopT1-1 and HopT1-1m3 and streptavidin-associated peptides containing each GW and GF motifs of HopT1-1 and HopT1-1m3**

**(A)** To perform the pull-down experiment, we first assessed the accumulation level of streptavidin-associated peptides containing each GW and GF motifs of HopT1-1 and HopT1-1m3 respectively, by using dot blot assay. The biotinylated peptides were immobilized with HRP-streptavidin beads then were spotted on a nitrocellulose membrane at three different amounts (1 µg, 0.1 µg and 0.01 µg). The presence of peptides was revealed by adding ECL substrate. **(B)** Agrobacterium tumefaciens strains carrying Myc-HopT1-1 or Myc-HopT1-1m3 constructs were infiltrated in four-week-old N. benthamiana leaves. Non-infiltrated N. benthamiana leaves were used as a control. Protein accumulation level of HopT1-1 and of HopT1-1m3 was monitored at 3 dpi by immunoblotting. Arrow indicates the band corresponding specifically to HopT1-1 and HopT1-1m3 detected using anti-Myc antibody. Relative quantification was performed using the unspecific proteins (*) detected by anti-Myc antibody. Both the proteins exhibit stable accumulation in planta. **(C)** To perform FRET-FLIM analysis, four-week-old N. benthamiana leaves were infiltrated with CFP-AGO1 alone or with YFP-tagged HopT1-1 or HopT1-1m3. After 2 dpi, protein accumulation level was monitored by immunoblotting using anti-GFP antibody. Ponceau staining was used to show equal protein loading for each sample. **(D)** Same as (C), but using CFP-AGO1 alone or with HopT1-1-HA. Nb: FRET occurs when the fluorescence of an acceptor molecule increases because of non-radiative transfer of energy from a donor molecule to the acceptor. However, quantitative interpretation of intensity-based images is sometimes difficult because of the number of factors that may contribute to intensity variations. For example, fluorophore intensity is related to both its local concentration and quantum yield (Gryczynski, Z., Gryczynski, I. & Lakowicz, J.R. in Molecular Imaging 21–56 Academic Press, 2005). To overcome such difficulties, FLIM (fluorescence lifetime imaging) has been developed as a complementary approach. Since lifetime is known to be related only to the fluorescence quantum yield; and not to the fluorescence intensity, lifetime measurements are not influenced by the accumulation level of the fluorescent fusion proteins. Therefore, the lower expression of HopT1-1m3 compared to HopT1-1 reported in this supplemental figure has no influence on our FRET/FLIM data showing that HopT1-1, but not HopT1-1m3, physically interact with AGO1. **(E)** HopT1-1 was manually docked through GW3 motif (W181) to arabidopsis AGO1 Trp-pocket-1 (left panel) and Trp-pocket-3 (right panel). Close-up views of W181 overlayed with AGO1 Trp-pocket-1 and Trp-pocket-3. HopT1-1 and AGO1 are shown in blue and grey, respectively and tryptophan residues are shown in orange. Extensive structural overlaps between AGO1 and HopT1-1 are highlighted with blue circles.
