## Supplemental Figure 4 for "A bacterial effector directly targets Arabidopsis Argonaute 1 to suppress Pattern-triggered immunity and cause disease"

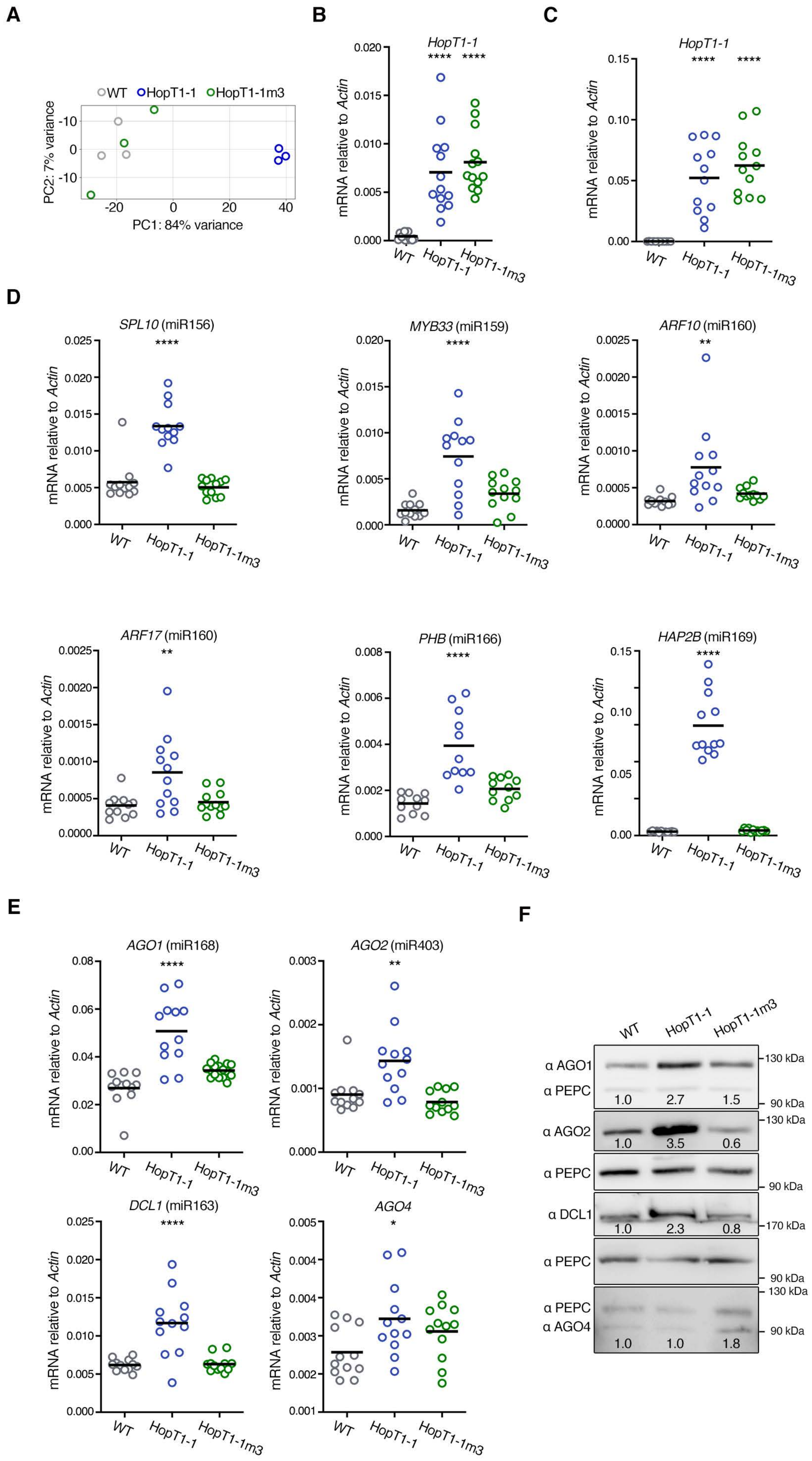
A
B
C
D
E
F
WT
HopT1-1
HopT1-1m3
PC2: 7% variance
PC1: 84% variance
HopT1-1
mRNA relative to Actin
SPL10 (miR156)
MYB33 (miR159)
ARF10 (miR160)
ARF17 (miR160)
PHB (miR166)
HAP2B (miR169)
AGO1 (miR168)
AGO2 (miR403)
DCL1 (miR163)
AGO4
α AGO1
α PEPC
α AGO2
α DCL1
α AGO4
130 kDa
90 kDa
170 kDa
1.0
2.7
1.5
3.5
0.6
2.3
0.8
1.8

**Supplemental Figure 4. HopT1-1 suppresses AGO1-mediated miRNA function in a GW-dependent manner**

**(A)** Principal component analysis based on gene expression RNA-seq data (log2 RPM) grouped according to treatment, each point represents a replicate. **(B)** The expression of HopT1-1 from the individuals described in Fig. 5B was monitored by RT-qPCR analysis in primary transformants expressing HopT1-1 or HopT1-1m3 under the constitutive 35S promoter. Number of individuals analysed in each condition is n=13. Actin was used as a control. **(C)** Biological replicate of the plants described in (B). The expression of HopT1-1 was monitored by RT-qPCR analysis in individual primary transformants expressing HopT1-1 or HopT1-1m3 under the constitutive 35S promoter. Number of individuals analysed in each condition is n=12. Actin was used as a control. **(D)** Expression of endogenous miRNA targets was monitored by RT-qPCR analysis in the same HopT1-1 transgenic plants depicted un (C) as well as in individual WT plants. The miRNA target transcripts that are analysed are indicated between brackets. Actin was used as a control. **(E)** Relative mRNA accumulation levels of RNA silencing factors targeted by miRNAs were monitored by RT-qPCR analysis in the same plants described in (C). AGO4 was used as an internal control, as this silencing factor is not targeted by any miRNA. Actin was used as a control. **(F)** Individuals of the plants depicted in (C) were pooled to monitor by immunoblotting the protein accumulation levels of the RNA silencing factors depicted in (D). PEPC protein accumulation level was used as loading control for each blot. Relative quantification of the protein accumulation using PEPC accumulation level was done using ImageJ software and is indicated below each condition. For RT-qPCR analyses, statistical significance was assessed by comparing the mean (black bar) of each condition with the mean of WT condition, using one -way ANOVA analysis (*: $p$-value < 0.05; **: $p$-value < 0.01; ****: $p$-value < 0.0001).
