## Supplemental Figure 5 for "A bacterial effector directly targets Arabidopsis Argonaute 1 to suppress Pattern-triggered immunity and cause disease"

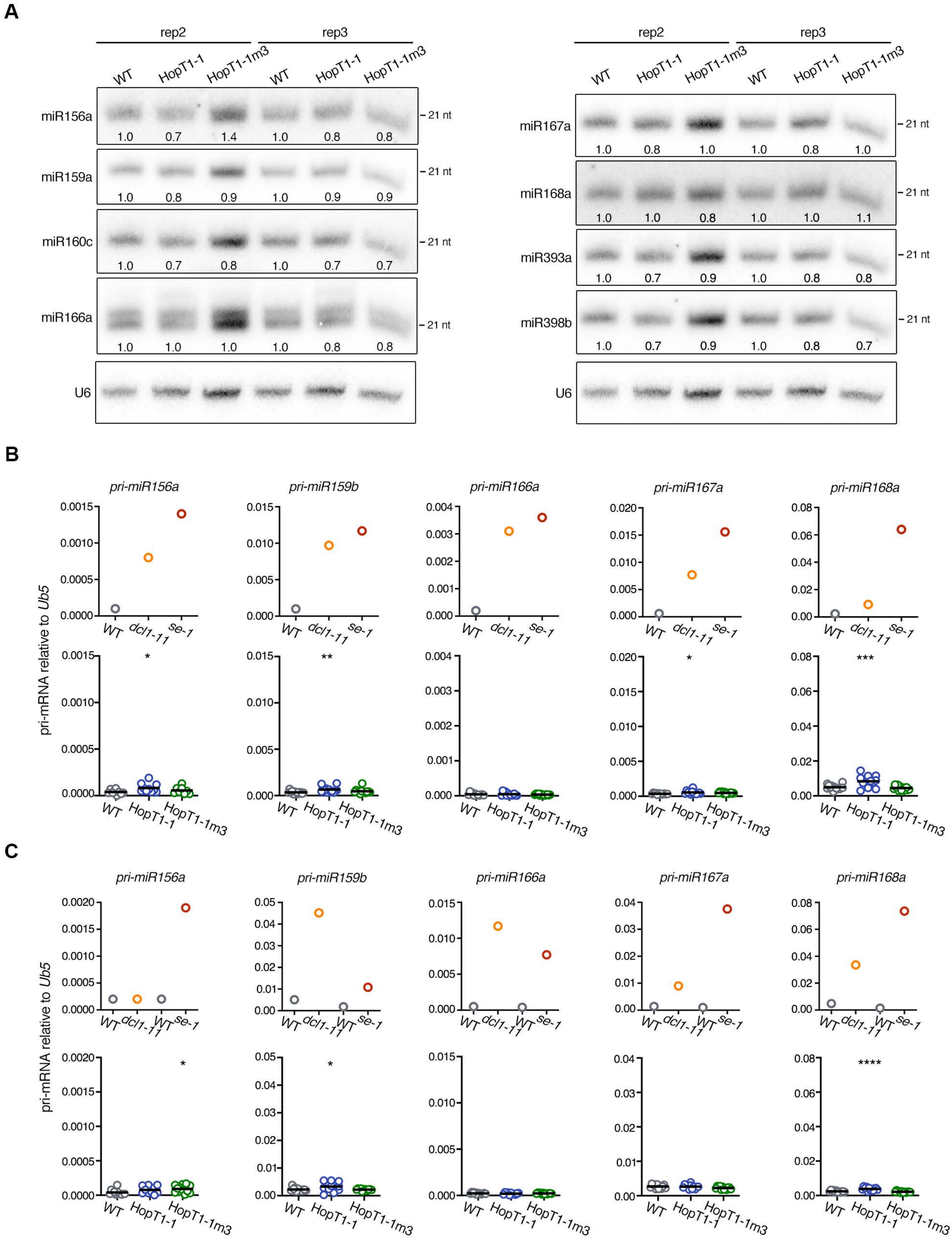


**Supplemental Figure 5. HopT1-1 does not substantially alter the accumulation of conserved Arabidopsis miRNAs nor of cognate pri-miRNAs**
**(A)** Accumulation levels of endogenous miRNAs in pooled WT plants and in pooled primary transformants expressing HopT1-1 or HopT1-1m3 were assessed by northern blot. U6 was used as a loading control. **(B)** Accumulation levels of endogenous pri-miRNAs were assessed by RT-qPCR in WT plants and in dcl1-11 and se-1 mutants (top panels) or in the above genotypes (bottom panels). Ubiquitin5 (ub5) was used as a control. For the RT-qPCR analysis of HopT1-1 and HopT1-1m3 transgenic plants, statistical significance was assessed by comparing the mean (black bar) of each condition with the mean of WT condition, using one-way ANOVA analysis (*: p-value<0.05; ****: p-value<0.0001). **(C)** Same than in (B) but for a biological replicate.
