## Supplemental Figure 6 for "A bacterial effector directly targets Arabidopsis Argonaute 1 to suppress Pattern-triggered immunity and cause disease"

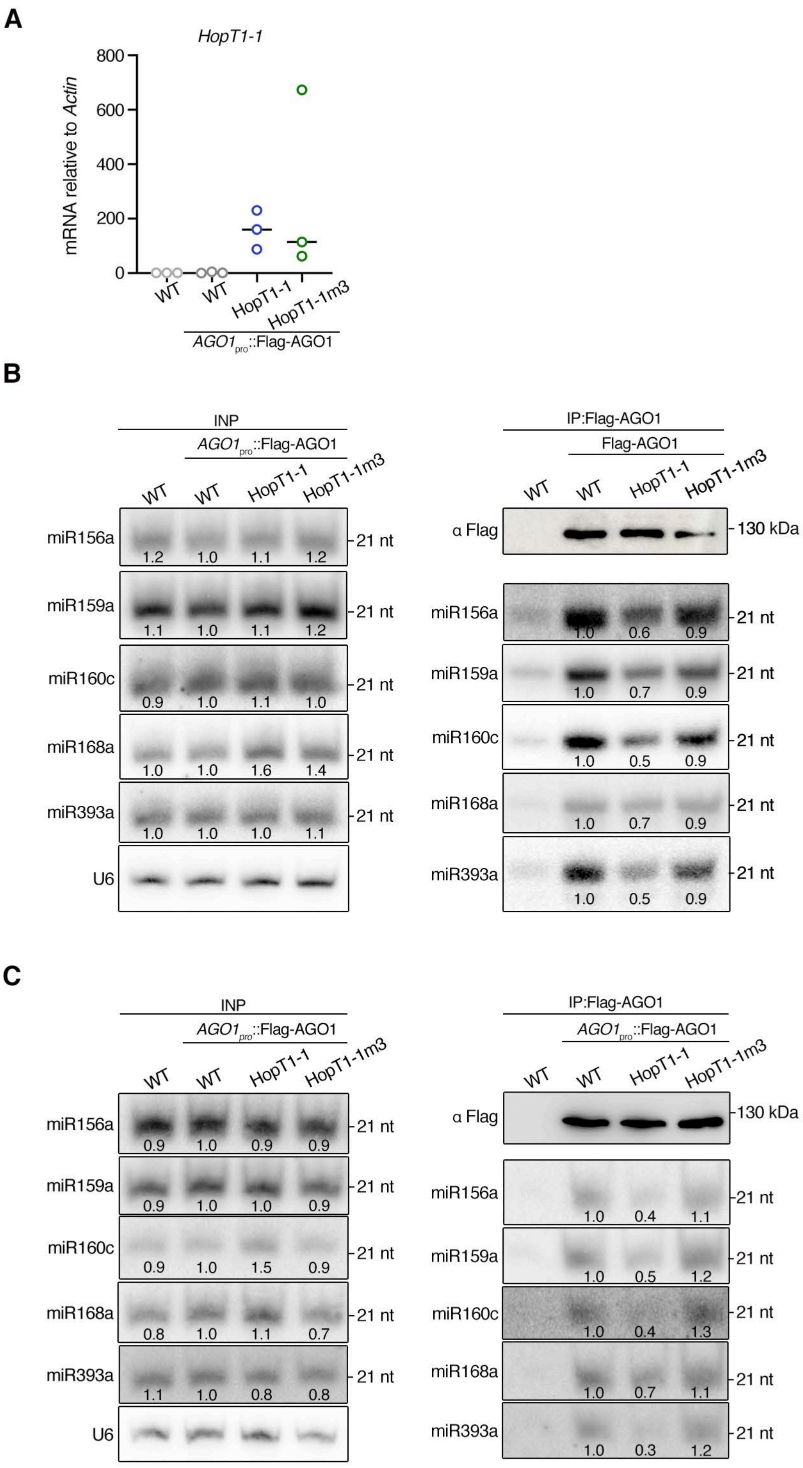


**Supplemental Figure 6. The AGO-binding platform of HopT1-1 interferes with the miRNA loading on AGO1**

**(A)** Inflorescences from WT plants or from AGO1pro::Flag-AGO1 transgenic line or from pooled primary transformants expressing HopT1-1 or HopT1-1m3 in AGO1pro::Flag-AGO1 transgenic plants were collected. Expression level of HopT1-1 transcript on these lines was assessed by RT-qPCR on three biological replicates. **(B)** On the tissues described in A, the accumulation level of canonical miRNAs was analysed by Northern blot with U6 probe used as a loading control (left panel). Flag-AGO1 was immunopurified from the inflorescence described in (A) and accumulation level of Flag-AGO1 proteins was assessed by western blot (right and upper panel). The accumulation level of miRNAs associated to Flag-AGO1 in each condition was analysed by Northern blot (right panel). **(C)** Same as (B) but for another biological replicate.
