## Supplemental Figure 7 for "A bacterial effector directly targets Arabidopsis Argonaute 1 to suppress Pattern-triggered immunity and cause disease"

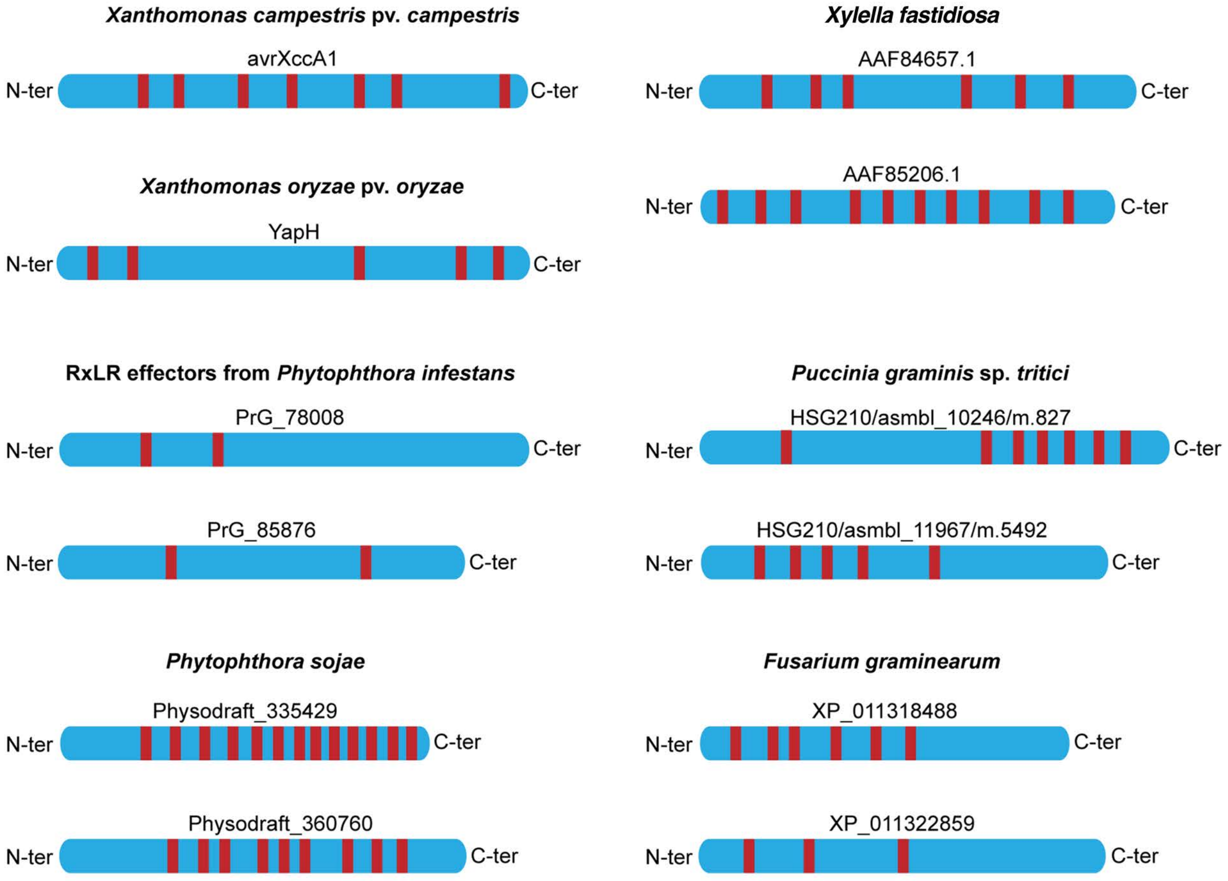


**Supplemental Figure 7. Several effectors encoded by agriculturally important phytopathogens contain canonical GW/WG motifs**
Effectors encoded by bacteria (*Xanthomonas campestris*, *Xanthomonas oryzae* and *Xyllela fastidiosa*), oomycetes (*Phytophthora infestans* and *Phytophthora sojae*) or fungi (*Puccinia graminis* and *Fusarium graminearum*) containing the highest score (matrix AGO-planVir) of GW/WG motifs prediction were retrieved by using the web portal http://www.comgen.pl/whub (Zielezinski A. & Karlowski WM, 2014). A red bar represents each GW or WG motif.
