## Supplemental Table 1 for "A bacterial effector directly targets Arabidopsis Argonaute 1 to suppress Pattern-triggered immunity and cause disease"

**Table S1, related to Experimental Procedures. Primers used in this study.**

| Name | Sequence | Remarks |
|---|---|---|
| PR1_F | 5'- AAAACTTAGCCTGGGGTAGCGG-3' | qPCR |
| PR1_R | 5'-CCACCATTGTTACACCTCACTTTG-3' | qPCR |
| PR2_F | 5'- GCTTCCTTCTTCAACCACACAGC-3' | qPCR |
| PR2_R | 5'-CGTTGATGTACCGGAATCTGAC-3' | qPCR |
| WRKY75_F | 5'-AGGCCGTCAAGAACAACAAG-3' | qPCR |
| WRKY75_R | 5'-ACGACCACTTCTTGGTCCAC-3' | qPCR |
| AGO1_F | 5'-AAGGAGGTCGAGGAGGGTATGG-3' | qPCR |
| AGO1_R | 5'-CAAATTGCTGAGCCAGAACAGTAGG-3' | qPCR |
| AGO2_F | 5'-GCCCCAATAACGCAGTTTTA-3' | qPCR |
| AGO2_R | 5'-CAAATTCGTTTCAACACACCA-3' | qPCR |
| AGO4_F | 5'-GCCATTTCTGTTGTTGCGCCGATC-3' | qPCR |
| AGO4_R | 5'-ACAGAAGAACATGGAGTTGGCGACGT-3' | qPCR |
| DCL1_F | 5'-GCACCGTTTGAAATACTTGAGG-3' | qPCR |
| DCL1_R | 5'-CGCTACTCCAACTTGAACACC-3' | qPCR |
| HAP2B_F | 5'-TGCTGCAATTTCAAAACCTG-3' | qPCR |
| HAP2B_R | 5'-GCCAAAGATGATTTGCCTGT-3' | qPCR |
| MYB33_F | 5'-GACATTCACCTGTTATGATT-3' | qPCR |
| MYB33_R | 5'-TGGAGACTGAATGTAAGTAT-3' | qPCR |
| HopT1_qPCR_F | 5'-GGCTAGCGAAAGTCGTGAAC-3' | qPCR |
| HopT1_qPCR_R | 5'-AACCCTTATCGAAGCCCACT-3' | qPCR |
| Ub10_F | 5'-GGCCTTGTATAATCCCTGATGAATAAG-3' | qPCR |
| Ub10_R | 5'-AAAGAGATAACAGGAACGGAAACATAGT-3' | qPCR |
| Ub_F | 5'-TGAAGTCGTGAGACAGCGTTG-3' | qPCR |
| Ub_R | 5'-GGGCTTCTCCATTGTTGGTC-3' | qPCR |
| Pri-miR156a_F | 5'-TCTCCCTCCCTCTCTTTGATTC-3' | qPCR |
| Pri-miR156a_R | 5'-CCCAACTCTTTCATTCACAATTAGT-3' | qPCR |
| Pri-miR168a _F | 5'-AGCCAAGTGATGTTGCCTTT-3' | qPCR |
| Pri-miR168a _R | 5'-TCATCATCGAAGCCTATCCACA-3' | qPCR |
| P4835 | 5'-CACCACCCTCTTACGGACAAGA-3' | deletion of *hopT1-1* |
| P4836 | 5'-GGGTATCGAGTGATTGCTGA-3' | deletion of *hopT1-1* |
| P4837 | 5'-CACCTCTCAAGGAAAGGCTTGAT-3' | deletion of *hopT1-1* |
| P4838 | 5'-GAAACGTTTGTCTCCGGCTA-3' | deletion of *hopT1-1* |
| P4839 | 5'-CACTTGAACGAGATCGCAGA-3' | deletion of *hopT1-1* |
| P4840 | 5'-GCATCAAGCCTTTCCTTGAG-3' | deletion of *hopT1-1* |
